# Different Purkinje cell pathologies cause specific patterns of progressive ataxia in mice

**DOI:** 10.1101/2023.08.29.555378

**Authors:** Dick Jaarsma, Maria B. Birkisdóttir, Randy van Vossen, Demi W.G.D. Oomen, Oussama Akhiyat, Wilbert P. Vermeij, Sebastiaan K.E. Koekkoek, Chris I. De Zeeuw, Laurens W.J. Bosman

**Author notes:** Correspondence to Dick Jaarsma or Laurens Bosman, PO Box 2040, 3000 CA Rotterdam, The Netherlands. The authors declare no competing interests.

## Abstract

**Background:** Gait ataxia is one of the most common and impactful consequences of cerebellar dysfunction. Purkinje cells, the sole output neurons of the cerebellar cortex, are often involved in the underlying pathology, but their specific functions during locomotor control in health and disease remain obfuscated.

**Objectives:** We aimed to describe the effect of gradual adult-onset Purkinje cell degeneration on gaiting patterns in mice and whether two different mechanisms that both lead to Purkinje cell degeneration caused different patterns in the development of gait ataxia.

**Methods:** Using the ErasmusLadder together with a newly developed limb detection algorithm and machine learning-based classification, we subjected mice to a physically challenging locomotor task with detailed analysis of single limb parameters, intralimb coordination and whole-body movement. We tested two Purkinje cell-specific mouse models, one involving stochastic cell death due to impaired DNA repair mechanisms (*Pcp2-Ercc1*^-/-^), the other carrying the mutation that causes spinocerebellar ataxia type 1 (*Pcp2-ATXN1[82Q]*).

**Results:** Both mouse models showed increasingly stronger gaiting deficits, but the sequence with which gaiting parameters deteriorated depended on the specific mutation.

**Conclusions:** Our longitudinal approach revealed that gradual loss of Purkinje cell function can lead to a complex pattern of loss of function over time, and this pattern depends on the specifics of the pathological mechanisms involved. We hypothesize that this variability will also be present in disease progression in patients, and our findings will facilitate the study of therapeutic interventions in mice, as very subtle changes in locomotor abilities can be quantified by our methods.

## Introduction

Gait ataxia, which is characterized by balance problems and walking abnormalities, is one of the more common consequences of cerebellar dysfunction, and is a key hallmark of a large number of acquired and inherited diseases (Earhart and Bastian, 2001; Stolze et al., 2002; Schmitz-Hübsch et al., 2006; Ilg et al., 2007; Morton and Bastian, 2007; Klockgether, 2010; Vinueza Veloz et al., 2015; Synofzik and Schule, 2017; Buckley et al., 2019; Buijsen et al., 2019; Klockgether et al., 2019; Lieto et al., 2019; Machado et al., 2020; Shah et al., 2021; Beaudin et al., 2022; Joyce et al., 2022; Castiglia et al., 2023; Coarelli et al., 2023). Clinical characteristics, severity and disease course vary between patients, and depend among others on the nature of the pathological mutation and the possible involvement of extracerebellar damage (van de Warrenburg et al., 2002; Klockgether et al., 2019; Diallo et al., 2020; Manto et al., 2020; Shah et al., 2021). Gait ataxia as a consequence of cerebellar degeneration is hardly ever an isolated feature, but occurs often together with problems with speech (Kent et al., 1979; Mariën et al., 2014), swallowing (Vogel et al., 2015; Sasegbon et al., 2020; Sasegbon and Hamdy, 2023) and breathing (Ebert et al., 1995; Sriranjini et al., 2010; Krohn et al., 2023), as well as with non-motor symptoms (Schmahmann, 1998; Hoche et al., 2018; Schmahmann et al., 2019; Malek et al., 2022). Effective treatments for most cerebellar disorders are currently lacking, although possible strategies are under intense investigation, with varying degrees of success (Buijsen et al., 2019; Mitoma et al., 2021; Bhartiya et al., 2022; Srinivasan et al., 2022; Synofzik et al., 2022).

Given the degenerative nature of many forms of cerebellar ataxia, potential treatments should preferably be aimed at pre-or prodromal stages of the disease. Careful observation of mutation carriers has revealed that abnormalities in the gaiting pattern can be detected at the prodromal stage, thus before clinical diagnosis (Jacobi et al., 2013; Rochester et al., 2014; Velázquez-Pérez et al., 2014a; Velázquez-Pérez et al., 2014b; Maas et al., 2015; Ilg et al., 2016; Velázquez-Pérez et al., 2017; Kim et al., 2021; Shah et al., 2021; Velázquez-Pérez et al., 2021; Thierfelder et al., 2022). Gait analysis is also central to the study of mouse models for cerebellar ataxia that play a crucial role in mechanistic and interventional studies to further our understanding of disease mechanisms, and to test and validate curative approaches for cerebellar ataxia (Cendelin et al., 2022).

A variety of behavioral paradigms has been used to detect gaiting deficiencies in mice, including footprint or video analyses of spontaneous locomotion, forced locomotion tests on treadmills, belts and rods, and skilled locomotor tests such as balance beam or horizontal ladder crossing tests (Jones and Roberts, 1968; Hartmann et al., 2004; Stanley et al., 2005; Zörner et al., 2010; Bellardita and Kiehn, 2015; Machado et al., 2015; Vinueza Veloz et al., 2015; Preisig et al., 2016; Darmohray et al., 2019; Timotius et al., 2019). These methods all show gaiting and/or balance deficits in mouse models of cerebellar disease, but vary in sensitivity and their ability to detect graded changes throughout disease progression (Hartmann et al., 2004; Stanley et al., 2005; Markvartová et al., 2010; Vinueza Veloz et al., 2015; Preisig et al., 2016). Tests that both capture early disease stages and monitor quantitative and qualitative changes associated with disease progression will be most effective in interventional studies.

In a previous study, where we tested a panel of cerebellar mutant mouse models on an automated horizontal ladder test (ErasmusLadder), we uncovered different profiles of quantitative and qualitative changes in locomotor parameters (Vinueza Veloz et al., 2015). The ErasmusLadder, which has an alternating pattern of high and low rungs and shelter boxes on either side, combines a challenging motor task with a clear motivation for the mice to engage, and a high number of repeats, allowing to differentiate between the performance of cell-specific cerebellar mutant mouse lines (Vinueza Veloz et al., 2015).

Following up on the demonstration that the ErasmusLadder enables the differentiation between cerebellar mutants in cross sectional studies (Van Der Giessen et al., 2008; Renier et al., 2010; Schonewille et al., 2011; van der Vaart et al., 2011; Baudouin et al., 2012; Saab et al., 2012; Vinueza Veloz et al., 2012; Galliano et al., 2013; Marques et al., 2015; Vinueza Veloz et al., 2015; Ha et al., 2016; Peter et al., 2016; Rahmati et al., 2016; French et al., 2018; Prekop et al., 2018; Sathyanesan et al., 2018; Sayed-Zahid et al., 2019; Wu et al., 2019; Almeida et al., 2020; Grasselli et al., 2020; Haify et al., 2020; Namdar et al., 2020; Peter et al., 2020; Blot et al., 2021; Lang-Ouellette et al., 2021; Vacher et al., 2021; White et al., 2021; Kaiser et al., 2022; Klomp et al., 2022; Lauffer et al., 2022; Ottenhoff et al., 2022; Birkisdóttir et al., 2023a; Fang et al., 2023) (Table S1), in this study we aimed to probe the ErasmusLadder as a tool for quantifying disease onset and progression in cerebellar disease models. We examined two mouse lines displaying progressive Purkinje cell degeneration, and followed their behavioral performance from early to severe symptomatic stages.

In one line, the loss of ERCC1/XPF nuclease function compromised several DNA repair mechanisms of Purkinje cells, and these mice display a stochastic loss of Purkinje cells (Weeda et al., 1997; de Graaf et al., 2013; Vermeij et al., 2016; Apelt et al., 2021; Birkisdóttir et al., 2023b). In the other, the transgenic overexpression of human ATXN1-82Q resulted in prolonged periods with Purkinje cell pathology, with cell loss occurring at a relatively low rate (Burright et al., 1995; Clark et al., 1997; Kaemmerer et al., 2001; White et al., 2021). In the mouse line with stochastic Purkinje cell loss, ataxia started mainly with a slower, less regular gait with smaller step, while in the mouse line with prolonged Purkinje cell pathology, ataxia started predominantly with a loss of coordination. Despite following different trajectories, eventually both mouse lines converged in their behavior, reaching a similar degree of ataxic gait. We conclude that quantitative gait analysis on the ErasmusLadder provides insight into different Purkinje cell-specific mechanisms underlying the control of locomotion in health and disease, and provides a tool for refined quantitative analysis of the effects of intervention.

## Methods

### Ethics Statements

Animal experiments were performed according to institutional guidelines as overseen by the Animal Welfare Board of the Erasmus MC, following Dutch and EU legislation. Prior to the start of the experiments, a project license for the animal experiments performed for this study was obtained from the Dutch national authority and filed under no. AVD101002015273.

### Mice

Pcp2-cre/*Ercc1*^*-/f*^ mice (designated PC-Ercc1 KO, hereafter) that are deficient for ERCC1 specifically in Purkinje cells were generated as described previously (de Graaf et al., 2013; Van’t Sant et al., 2021; Birkisdóttir et al., 2023b). First, *Pcp2-Cre*^*+*^*/Ercc1*^*-/+*^ mice on a C57BL/6J background were generated by crossing female Tg(Pcp2-Cre)Mpin mice (Barski et al., 2000) with male *Ercc1*^*-/+*^ mice. Subsequently, female *Pcp2-Cre*^*+*^*/Ercc1*^*-/+*^ were crossed with male *Ercc1*^*f/f*^ mice in FVB background to yield *Pcp2-Cre*^*+*^*/Ercc1*^*-/f*^ in a uniform C57BL/6JxFVB F1 hybrid background. *Pcp2-Cre*^*+*^*/Ercc1*^*+/f*^ and *Pcp2-Cre*^*-*^*/Ercc1*^*-/f*^ mutant mice that do not develop a degenerative phenotype served as controls (de Graaf et al., 2013; Van’t Sant et al., 2021).

Transgenic mice that overexpress human *ATXN1* cDNA with an 82 CAG repeat under the Pcp2-promoter (Tg(Pcp2-ATXN1*82Q)5Horr) were bred from strain “B05” as described in Burright et al. (1995), and we refer to these mice as PC-Sca1 in this report. For this strain, heterozygous transgenic mice were compared with WT littermates. These mice were kept on an FVB background, and fed with standard diet.

Animals were group housed in a general mouse facility. Environment was controlled with a temperature of 20–22°C and 12 h light:12 h dark cycles. Food and water were offered ad libitum. Mice were weighed weekly, and scored blindly for gross morphological and motor abnormalities daily.

### Analysis of Purkinje cell survival

Histological procedures and stereological quantification of Purkinje cells were done as previously described (Van’t Sant et al., 2021; Birkisdóttir et al., 2022). Staining was performed on 40 μm-thick serial transverse freezing microtome sections of gelatin-embedded 4% paraformaldehyde transcardially perfused cerebella (Van’t Sant et al., 2021). Sections were processed free-floating for immunofluorescence (Jaarsma et al., 2018). Primary antibodies (supplier; catalogue number; RRID number; dilution) used in this study were as follows: rabbit anti-Calbindin (Swant; C9638; AB_2314070; 1:20,000), mouse anti-Calbindin (Sigma Aldrich; C8666; AB_2313712; 1:20,000), goat anti-FOXP2 (Santa Cruz Biotechnology; sc21069, AB_2107124; 1:1,000), mouse anti-NeuN, clone A60 (Millipore; MAB377; RRID:AB2298772; 1:4,000), guinea pig anti-VIAAT (Synaptic Systems; 131004; AB_887873; 1:1,000), and rabbit anti-VIAAT (Synaptic Systems; 131003; AB_887869; 1:1,000). Alexa488-, Cy3-, and Cy5-conjugated antibodies raised in donkey (Jackson ImmunoResearch) diluted at 1:200 were used as secondary antibodies for visualization. Immuno-fluorescence sections were analyzed and imaged using Zeiss LSM700 and Leica Stellaris 5 confocal microscopes, and Zeiss Axio Imager D2 widefield microscopes.

For stereological quantification of Purkinje cells, we used serial transverse sections double-immunostained for Calbindin and FOXP2. Sections were analyzed using the Optical Fractionator tool of a StereoInvestigator software package (MBF Bioscience), integrated in a Zeiss LSM700 confocal microscope setup as recently described (Birkisdóttir et al., 2021; Birkisdóttir et al., 2023b).

### Accelerating rotarod

In the rotarod test (Ugo Basile, Varese, Italy), animals had to walk on a cylinder with a diameter of 3 cm that gradually accelerated from 2 to 40 or to 80 rotations per minute (rpm) during 300 s, respectively. Performance was assessed by measuring the time spent on the rotarod until the mice fell off the cylinder, with 300 s as the maximum time. Training consisted of four trials (with 1 h intervals) on the 2-40 rpm protocol for first days, followed by 2 trials at 2-40 rpm and 2-80 rpm on the consecutive day. Weekly testing consisted of two trials with an interval of 1 h at 2-40 rpm, followed by two trials at 2-80 rpm.

### ErasmusLadder

The ErasmusLadder (Noldus, Wageningen, The Netherlands) consists of a horizontal ladder counting 37 alternating low and high rungs on each side in between two shelter boxes (Vinueza Veloz et al., 2015). Each rung of the ErasmusLadder is connected to a touch sensor, and sensor output is stored together with meta-data in a MySQL database.

After one session in which the mice were allowed to explore the ErasmusLadder freely without any incentives to walk, the mice were trained during five daily sessions of 42 crossings each to walk between the two shelter boxes. Mice are motivated to leave the boxes by a strong tail wind that was preceded by a three second LED light stimulus (Fig. S1A). Escapes from the start box before the light was turned on (early escapes) triggered a strong head wind inducing the mouse to return to the start box to start a new trial. Trials in which the mice returned to the starting box (irregular trials) were also discarded; the mouse having to redo the trial (Fig. S1A). After the five daily training sessions in the first week, mice were tested once a week during 14 weeks. The test periods were based on rotarod data, and were from 13 to 26 weeks for PC-Ercc1 KO and 7 to 20 weeks for PC-Sca1 mice.

### Gait analysis

Instead of using the analysis software from the manufacturer, we retrieved the raw data directly from the MySQL database, and employed refined filter settings and analysis algorithms, using custom software written in Python. First, we removed data from irregular trials and deviant rung touches using the following steps: exceptionally long trials (>15 s) resulting from pausing animals, were excluded from further analysis; this affected <1% of all trials. Second, we disregarded abnormally short touches (<35 ms) resulting from rung vibration after release, as well as abnormally long touches (> 800 ms), well above the duration of regular touches. Third, we excluded the first (1-6) and last rungs (31-37) from the analysis as these rungs relate to stepping out of the start box, and into the end box, respectively. Finally, we eliminated trials with extraordinary numbers of rung touches, excluding trials with < 5 or > 150 registered touches.

Following the filtering steps, we associated touches with limbs. This step was based on the assumption that limbs are only placed at the ipsilateral side of the ErasmusLadder, so right limbs only touch the rungs of the right side, and left limbs only those on the left. We also assumed that mice would always exit the start box with their forelimbs first, so that the first touch on the right corresponds to the right forelimb, and the first touch on the left to the left forelimb. From this, we attributed every consecutive touch to one of the limbs, taking into account that a new touch could only be made by a limb that did not already touch a rung.

After this iteration, a number of touches could not directly be attributed to one of the limbs. The algorithm tried subsequently to predict the correct limb for each of the unassigned touches. To this end, it calculated the probability of each limb to reach the rung on which a touch was assigned, based upon the distance (<7 rungs per step) and the required swing time (>15 ms). If no limb could be associated to a specific touch, the algorithm went one step back and tried with another fit to the previous step to obtain a result. If this also failed, we considered the option that a mouse had both front and hindlimbs simultaneously on the same rung, but it only for touch duration that >150% of the average touch duration of that rung. If splitting a single touch in a forelimb and a hindlimb touch resulted in a valid pattern, we used these values. If none of these helped, the touch was registered as undetermined. Overall, around 95% of the touches could be classified in this way. Unclassified touches were not included in the further analysis.

After establishing a possible gaiting pattern for a trial, we consecutively tested the validity of this pattern, and rejected the whole trial from further analysis when a limb is attributed to more than one rung at a time, or when a gap in the coverage is detected. The latter implies that a mouse appears to make a step with a length of 8 or more rungs.

### Validation of the algorithm

To verify the output of the automated limb detection, we made video recordings (200 frames/s) of two adult control mice with a C57BL/6J background. Manual analysis of these videos identified 345 steps during 43 trials. Of these, 322 (93%) were correctly classified. The steps that were not classified correctly mostly involved slips or timing errors. In the latter case, the ErasmusLadder reported much shorter touches than seen on the video. As the moment of activation of the pressure sensor cannot be seen on the video, it can be that mice were in contact with a rung for a longer time than registered by the pressure sensor. It should be noted that chaotic gaiting patterns are associated with more errors in limb detection.

### Visualization and quantification of gaiting patterns

To visualize gaiting patterns on the ErasmusLadder, we first created gait diagrams in which we color-coded the stance phase of each limb, i.e., the time when each limb touched a rung, as opposed to the swing phase. For each combination of limbs, we calculated the fraction of each trial in which they were simultaneously in stance phase. Next, we focused on the diagonal stance, and performed a linear phase transform of the step cycle of the right forelimb, with 0 = start of stance and *π* = start of swing phase. In this step cycle of the right forelimb, we plotted the start of the stance phase of the left hindlimb. The circularity of the obtained graph with 0.1 *π* bins was calculated as 4*π* x area / perimeter^2. The eccentricity was determined by calculating the center of gravity of the same shape and then calculating the distance of the center of gravity to the origin.

For each mouse line and stance (diagonal, lateral and girdle), we also created two-dimensional Hildebrand plots (Hildebrand, 1965; Cartmill et al., 2020). On the x-as, we plotted the fraction of the step cycle during which the right forelimb touched a rung, and on the y-as the fraction of the step cycle between the placement of the right forelimb and a second limb, e.g., the left hindlimb for the diagonal stance. The density of the events was normalized per session, and, for each mouse line and stance, we used unsupervised machine learning in combination with *k* means clustering to divide the histograms in two clusters.

To extract image features from the two-dimensional histograms (stored as TIFF-images), we used the pretrained InceptionV3 unsupervised machine learning algorithm from the Keras applications library in Python, whereby we removed the output layer of the pretrained model. The extracted image features were used as input for *k* means clustering (*k* = 2). To control for model sensitivity to the initial conditions, we ran each model ten times and compared the resultant clusters.

### Principal component analysis

For each session of the two mouse lines we used 23 different parameters to perform principal component analysis (PCA), i.e., run time, number of steps, number of even steps, number of steps with step size = 2 rungs, number of steps with step size = 4 rungs, number of steps with step size = 6 rungs, average stance phase of each of the paws, number of high rungs touched by forelimbs and hindlimbs, number of lower rungs touched by front and hindlimbs, median duty cycle (= stance phase) of the right front paw, median CV2 of the step times of the right front paw, percentages of time where the mice has either a diagonal, girdle, lateral, 1 paw, 3 paw or 4 paw stance. All of these parameters refer to individual runs/trials. To perform the PCA, the Scikit-learn Python package was used. the PCA was instructed to retain as much principal components as needed to explain at least 85% of the total variance in our datasets.

### Statistical analysis

Unless stated otherwise, for each session, data for control and mutant mice were compared using a Mann-Whitney test, followed by Benjamini-Hochberg correction for multiple comparisons. Only *p* values that were found to be statistically significant after correction were reported as such. See Tables S2-S12.

## Data Sharing

Data and code generated for this manuscript will be made available in a public depository upon publication.

## Results

### Progressive adult-onset Purkinje cell degeneration in Purkinje cell-specific ERCC1-deficient mice is associated with progressive changes in walking speed and stepping patterns

To explore whether the ErasmusLadder can be used to precisely monitor progression of gait abnormalities over time in mouse models of cerebellar disease, we employed two mouse models with progressive Purkinje cell degeneration. In both lines, Purkinje cells were largely preserved until early-adult age. The first model, the PC-Ercc1 KO mouse, shows a complete loss of ERCC1/XPF nuclease function in post-natal Purkinje cells, affecting several DNA repair pathways, in particular nucleotide excision repair and inter-strand crosslink repair (Weeda et al., 1997; de Graaf et al., 2013; Vermeij et al., 2016; Apelt et al., 2021; Birkisdóttir et al., 2023b). A previous proteome study indicated that PC-Ercc1 KO mice show largely preserved Purkinje cells at 8 weeks, and moderate to severe loss of Purkinje cells at 16 and 26 weeks, respectively (de Graaf et al., 2013).

In a cohort that we analyzed for loss of locomotor function using the accelerating rotarod test, we found that PC-Ercc1 KO mice had similar weights as control littermates (Fig. 1A), and developed a trend of reduced rotarod performance from 20-22 weeks of age, and significant deficits from 24 weeks of age (at 24 weeks of age, control mice: 281 [68] s; PC-Ercc1 KO mice: 175 [39] s; medians [inter-quartile range]; *p* = 0.006, Mann-Whitney test, statistically significant after Benjamini-Hochberg correction for multiple comparisons; see Table S2 for more detailed statistics; Fig. 1B). Performance on a more challenging version of the accelerating rotarod did not reach statistical significance at any age tested (Fig. 1C).

**Figure 1.**
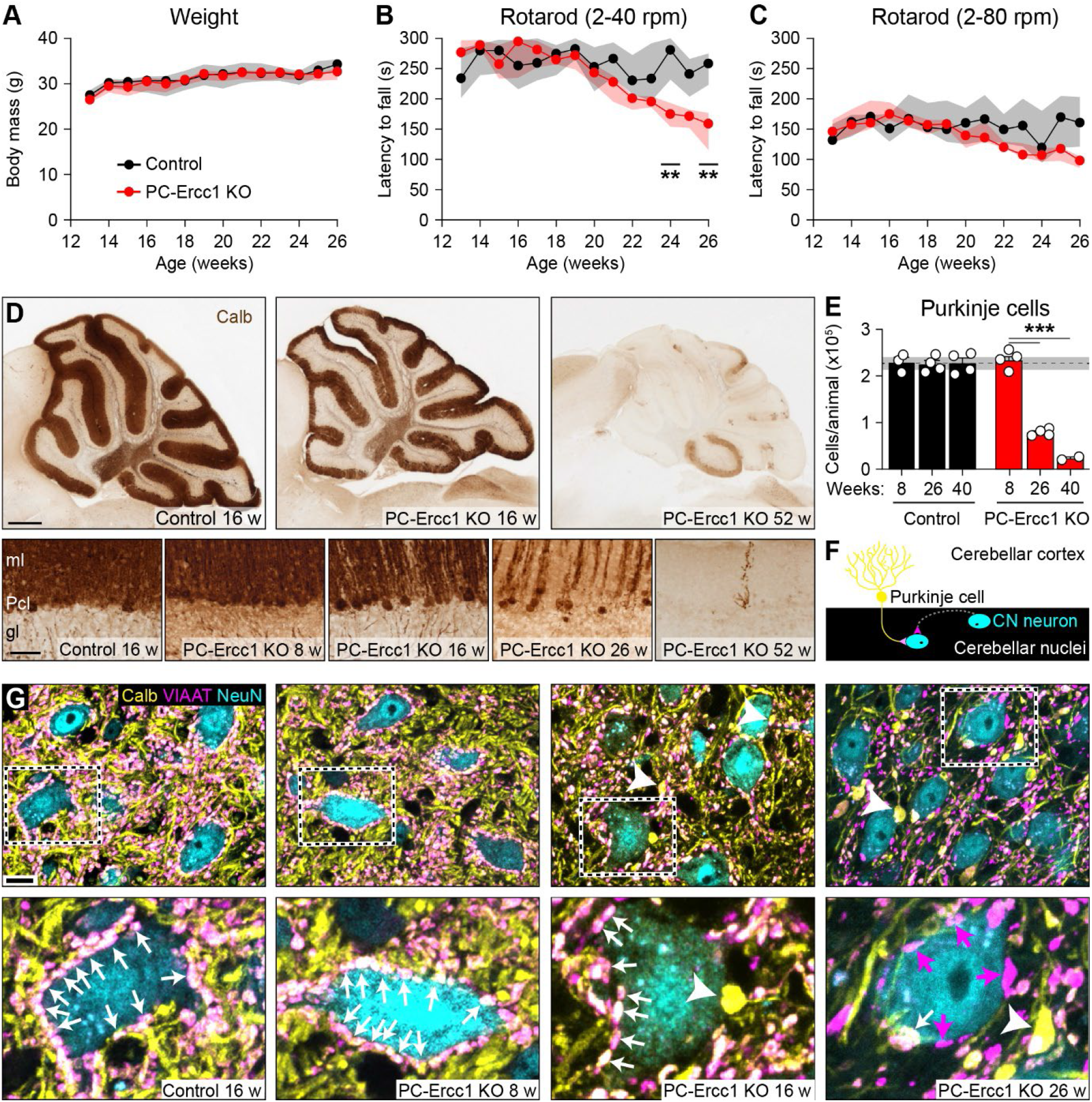
Purkinje cell-specific DNA repair-deficient mice develop adult-onset Purkinje cell degeneration. Body weight (**A**) and accelerating rotarod performance in PC-Ercc1 KO mice over time. PC-Ercc1 KO mice develop reduced performance, which was more pronounced at slow (2-40 rpm, **B**) than at fast acceleration (2-80 rpm, **C**). A-C: Medians with interquartile ranges; *n* = 8 male mice per group. **D**. Calbindin immunoperoxidase-DAB histochemistry illustrating loss of Purkinje cells in PC-Ercc1 KO mice starting after 8 weeks of age. Note the severe loss and near-complete absence of calbindin staining at 26 and 52 weeks of age, respectively. gl, granular layer; ml, molecular layer; Pcl, Purkinje cell layer. **E**. Stereological quantification of surviving Purkinje cells, showing severe loss in PC-Ercc1 KO mice at 26 and 40 weeks. **F**. Color scheme for G: yellow (calbindin) represents Purkinje cell axons and terminals, magenta (VIAAT) inhibitory nerve terminals (hence, Purkinje cell terminals are light magenta, whereas terminals of inhibitory interneurons are dark magenta), and cyan (NeuN) neuronal cell bodies. **G**. Confocal images of the lateral cerebellar nucleus. Dashed areas are enlarged in the lower row. Light magenta arrows point to exemplary Purkinje cell axon terminals (double-labelled for calbindin and VIAAT) that synapse on neuronal cell bodies. Note the reduced numbers and near-absence of Purkinje cell axon terminals, as well as Purkinje cell axonal swellings and swollen axon terminals (white arrow heads) in PC-Ercc1 KO mice at 16 and 26 weeks. At 26 weeks, numerous calbindin-negative VIAAT-positive boutons (representing non-Purkinje cells inhibitory nerve terminals; white arrows) are visible in PC-KO mice. Scale bars: 500 and 50 µm (D), and 10 µm (G). ** *p* < 0.01, Mann-Whitney test with Benjamini-Hochberg correction, *** *p* < 0.001, Tukey’s multiple comparison post-test.

Pathological analysis of Purkinje cell degeneration using calbindin immunohistology showed that at 8 weeks of age, the morphology and total number of Purkinje cells in PC-Ercc1 KO mice is similar to those in control mice. From 16 to 52 weeks, PC-Ercc1 KO mice showed mild to near-complete loss of Purkinje cells (Fig. 1D), with less than 30 and 10% residual Purkinje cells at 26 and 40 weeks, respectively (Fig. 1E). Calbindin staining further revealed largely normal Purkinje cell axons and terminals in the cerebellar nuclei at 8 weeks, a moderate reduction at 16 weeks, and a strong reduction at 26 weeks in PC-Ercc1 KO mice (Fig. 1F, G). The latter was accompanied by abnormal swellings of the axons and the terminals in the cerebellar nuclei (Fig. 1G). The loss of Purkinje cell terminals was confirmed by double staining for calbindin and vesicular inhibitory amino acid transporter (VIAAT, also termed VGAT or SLC32A1) to outline Purkinje cell presynaptic terminals on cerebellar nuclear neurons at 16 and 26 weeks (Fig. 1G). Interestingly, at 26 weeks the majority of VIAAT-positive boutons are calbindin-negative, reflecting non-Purkinje cell inhibitory synapses, which is consistent with findings in early onset Purkinje cell degeneration mouse models, such as Lurcher and Purkinje cell degeneration (Pcd) mice (Grüsser-Cornehls and Bäurle, 2001; Sultan et al., 2002).

To test the performance of PC-Ercc1 KO mice on the Erasmus Ladder, we used a protocol based on that of our previous study (Vinueza Veloz et al., 2015), but with a number of modifications: each mouse had to perform 19 daily sessions, starting with 5 training sessions at 12 weeks of age (sessions 1-5) followed by one daily session per week in subsequent weeks (Fig. 2A). During each session, each mouse had to perform 42 trials, i.e., ladder crossings between the shelter boxes. Mice were motivated to leave the shelter boxes by a visual stimulus followed by a strong tail wind. Trials in which a mouse left the start box before the onset of the light (early escape), or returned to the starting box (irregular trials) were discarded, the mouse having to redo the trial (Fig. S1A). Box exit behavior was largely similar between PC-Ercc1 KO and control mice throughout the experiments (Fig. S1B). Trial times were similar between PC-Ercc1 KO and control mice at young ages. However, PC-Ercc1 KO mice showed increased trial times after 18 weeks of age (Fig. S1C).

**Figure 2.**
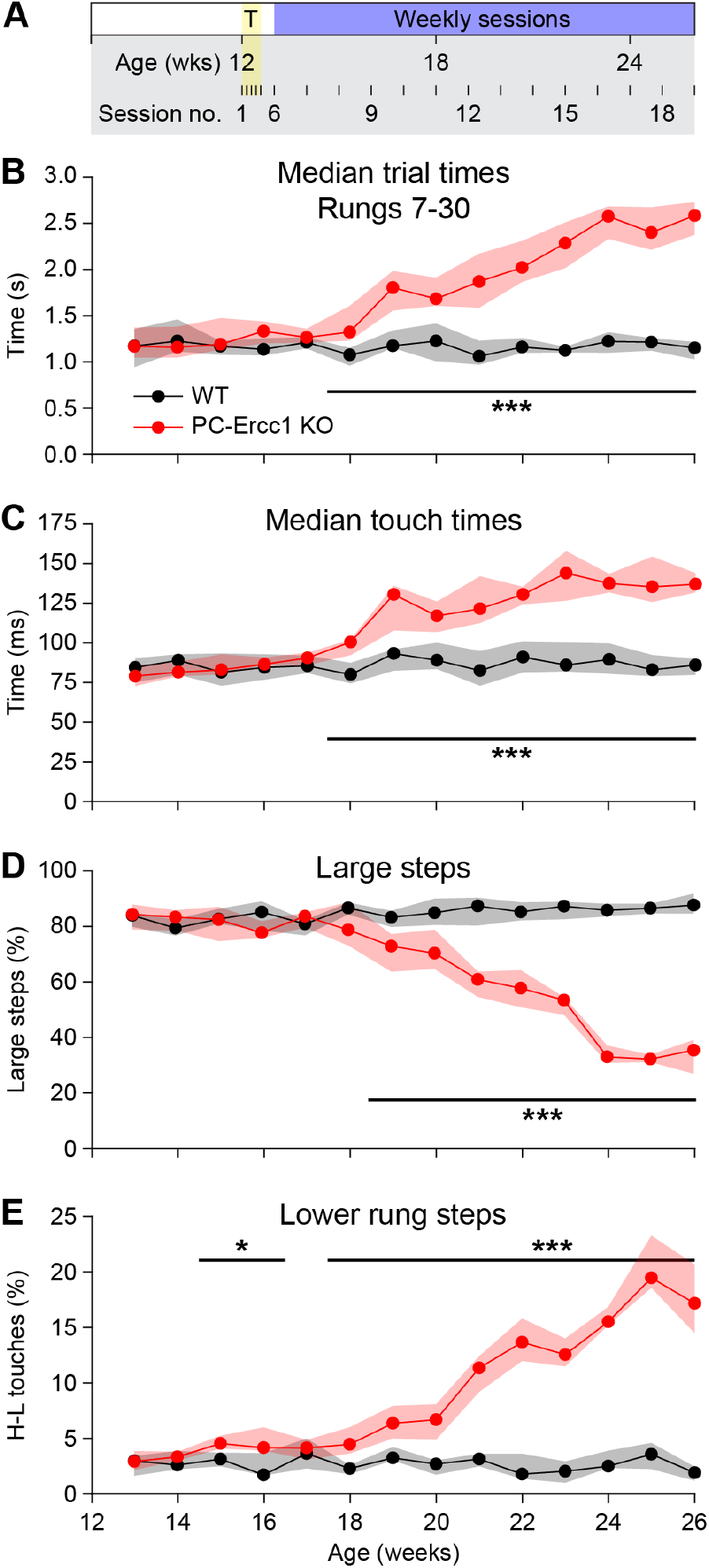
Purkinje cell degeneration is associated with altered stepping behavior A. At 12 weeks of age, the PC-Ercc1 KO mice and their controls were trained with five daily sessions on the ErasmusLadder, while from 13 weeks of age on they were tested weekly. **B**. Trial times. **C**. Touch times. **D**. Percentage of steps to a higher rung with a step size of 4 or more rungs. **E**. Percentage of steps from higher to lower rungs. Median values and interquartile ranges. * *p* < 0.05, *** *p* < 0.001, Mann-Whitney tests with Benjamini-Hochberg correction.

Gait and locomotion analysis with the ErasmusLadder is based on measuring rung touches by pressure sensors coupled to the rungs. In this study we used an updated algorithm (see Methods) that calculates the most likely pattern of limb placement for all limbs (Fig. S1D), allowing a more in-depth analysis of gaiting compared to algorithms used in previous studies that only analyzed forelimb steps (Vinueza Veloz et al., 2015; Haify et al., 2020). Comparison with video analysis showed that 93% of sensor touches were interpreted correctly by the algorithm (see Methods). The new algorithm filters for interrupted trials with excessive pauses or extreme numbers of sensor touches resulting from mice pausing or making turns on the ladder, and, excludes rung touches from the initial and final rungs after the start box and before the end box, respectively. Importantly, while these filtering steps resulted in shorter observed median trials times, they did not affect the differences in trial times between control and mutant mice (compare Fig. 2B, Fig. S1C). Thus, also after filtering, PC-Ercc1 KO mice show significantly increased crossing times starting from 18 weeks of age, while control mice show similar trial times throughout all sessions (Fig. 2B). Significantly, analysis of individual mice indicated that changes in trial times were remarkably similar in PC-Ercc1 KO mice, all showing a gradual increase in trial time starting from 18 weeks (Fig. S2A-C).

To enable comparison with our previous study (Vinueza Veloz et al., 2015), we first analyzed forelimb steps only. Consistent with increased trial times, PC-Ercc1 KO mice from 18 weeks of age started to show altered stepping behavior characterized by increased median durations of touching the rungs (Fig. 2D, S2C). Simultaneously, the stepping pattern became more irregular in PC-Ercc1 KO mice (Fig. S2E), while the ratio between stance and swing phase remained largely constant (Fig. S2F). Wild-type and young PC-Ercc1 KO mice typically walk on the upper rungs and mostly make relatively long steps encompassing 4-6 rungs (Fig. 2D, S2G,H). After 19 weeks, PC-Ercc1 KO mice showed reduced frequency of long steps, and increased frequency of short steps encompassing 2 rungs (Fig. S2G-I). PC-Ercc1 KO mice also showed an increased frequency of steps on lower rungs (Fig. 2E, Fig. S2H), a feature consistently found in cerebellar mutants in previous studies that has been interpreted as missteps resulting from decreased motor coordination (Van Der Giessen et al., 2008; Vinueza Veloz et al., 2015; French et al., 2018; Sathyanesan et al., 2018; Almeida et al., 2020; Namdar et al., 2020; Peter et al., 2020; Riemslagh et al., 2021; Birkisdóttir et al., 2023b). Interestingly, stepping on lower rungs was already significantly increased at week 15, which is before the onset of reduced speed. Video analysis of some older PC-Ercc1 KO mice confirmed that lower rung touches of forelimbs were associated with inadequate limb positioning (video S1). Thus, forelimb analysis shows that PC-Ercc1 KO mice develop changes in a number of gaiting parameters mostly related to speed and step length, while the earliest changes were linked to lower rung touches.

### Reduced inter-limb coordination in PC-Ercc1 KO mice

We next performed gait analysis based on the placement of all limbs. As illustrated by the representative gaiting diagrams (Fig. 3A, Fig. S3A), gaiting of control mice is most frequently characterized by sequential lifting of a forelimb, followed by the diagonal hindlimb, the opposite forelimb, and then the ipsilateral hindlimb. Analysis of the frequency of stance patterns indicates that about half of the time the animals stand on the rungs with either 3 limbs or the diagonal pair of limbs (Fig. 3B). Gaiting diagrams of PC-Ercc1 KO mice at a stage with severe Purkinje cell loss, showed more irregular patterns compared to controls (Fig. 3A, Fig. S3A). Even at that age, PC-Ercc1 KO mice, however, did not show major differences in the relative occurrences of stance patterns (Fig. 3B). For instance, the percentage of time spent with two diagonal limbs simultaneously touching the rungs, did not differ between controls and PC-Ercc1 KO mice throughout the sessions, except for the last three weeks showing a trend of reduced percentage of two limb diagonal stance, but not reaching the threshold for statistical significance (Fig. 3C). As a second approach of four limb gait analysis, we performed phase analysis of diagonally positioned limbs, by normalizing the stance and swing phases of a forelimb to the step duration, and plotting the phases of placement of the contralateral hindlimb (Fig. S3B). Control mice typically showed two preferred phases for hindlimb placement in the first and second half of the stance phase, respectively, while these peaks were less consistent in PC-Ercc1 KO mice (Fig. S3B). Statistical evaluation of the phase relations between the placement of diagonal limbs revealed only a clear lack of phase preference (“circularity”) at the final week of testing (Fig. S3B2), while the preference of hindlimb placement during the stance phase of the right forelimb (“eccentricity”) was not consistently affected in PC-Ercc1 KO mice (Fig. S3B3).

**Figure 3.**
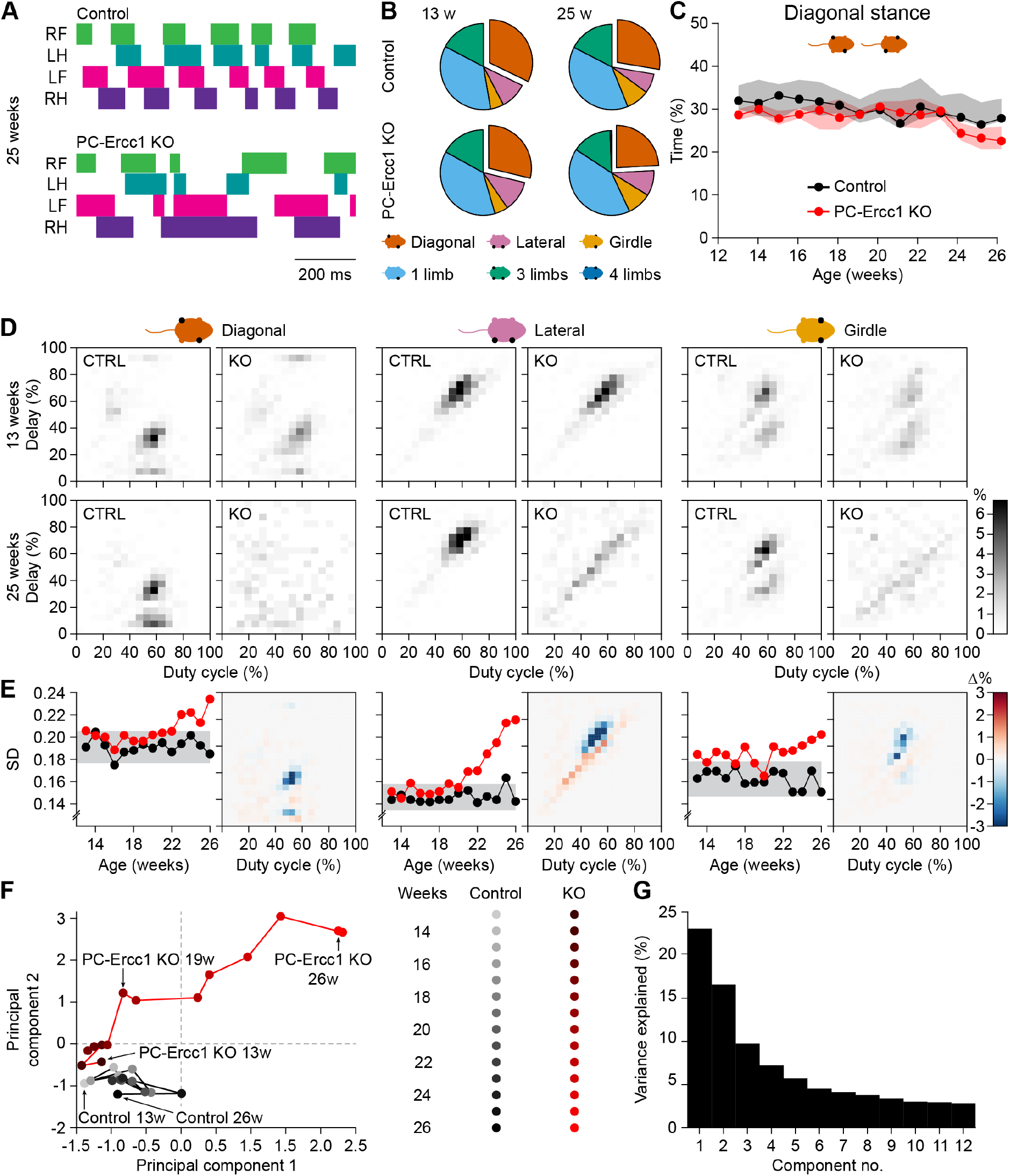
Reduced inter-limb coordination in PC-Ercc1 KO mice. **A**. Exemplary gaiting diagrams at 25 weeks of age, showing regular and irregular limb placement. The colored bars represent stance phase. **B**. Fraction of the touch times with one, two, three or four limb support. Two-limb support is split into diagonal, lateral and girdle stance. **C**. The occurrence of diagonal stance remained constant throughout time. **D**. Hildebrand plots representing the combinations of duty cycle (= percentage of the step cycle during which the right forelimb touches a rung) and the delay between placement of the right forelimb, and the left hindlimb (diagonal), right hindlimb (lateral), and left forelimb (girdle), respectively, at 13 and 25 weeks of age. **E**. Left panels: standard deviation of Hildebrand plots, with the 95% confidence intervals of control mice (grey bars). Right panels: differences between control and abnormal patterns as classified by a machine learning algorithm (see Fig. S4). **F**. Principal component analysis revealed that both groups of mice initially showed comparable behavior, that diverged later on. **G**. Variance explained for each principal component.

To further examine differences in inter-limb coordination between control and PC-Ercc1 KO mice, we generated two-dimensional plots based on the method of Hildebrand (Hildebrand, 1965), where the x-axis shows the duty cycle of a reference limb (here, the fraction of each step during which the right forelimb was touching a rung) and the y-axis shows the relative time of placement of a second limb (left hindlimb in diagonal stance; right hindlimb in lateral stance; and left forelimb in girdle; Fig. 3D, Fig. S4). This analysis revealed stereotyped patterns with one or few foci for each pair of limbs throughout all sessions in control mice. PC-Ercc1 KO mice showed the same patterns as controls at young age, but stereotypic pattern disappeared as Purkinje cell degeneration progressed (Fig. 3D, Fig. S4). To substantiate the changes in Hildebrand plots in PC-Ercc1 KO mice, we plotted the standard deviation of pixel intensities of each session. These plots showed the onset of altered inter-limb coordination in PC-Ercc1 KO mice to occur at 13, 21 and 23 weeks for girdle, lateral and diagonal limb pairs, respectively (Fig. 3E). Further analysis of Hildebrand plots using an unsupervised machine learning algorithm that grouped plots into either ‘control’ or ‘affected’ patterns, confirmed the analyses based on variation of pixel intensities: PC-Ercc1 KO mice show early onset of altered girdle stance, and later onset of altered diagonal and lateral stances (Fig. S4). Thus, Hildebrand plots expose changes in stereotypic inter-limb coordination following Purkinje cell degeneration Fig. 3D, E; Fig.S4).

To summarize the changes in gaiting pattern in PC-Ercc1 KO mice, we used principal component analysis on 23 gaiting parameters (see Methods). During early stages, the PC-Ercc1 KO mice were close to, but never overlapped with, control mice. From 19 weeks onward, the PC-Ercc1 KO mice started to more prominently diverge from the control mice, first along the second principal component, and subsequently also along the first (Fig. 3F-G). Thus, with the ErasmusLadder we could identify several gaiting parameters that deteriorated along with the loss of Purkinje cells in our PC-Ercc1 KO mice. Some gaiting parameters were already affected at early stages, while others came only later, and some were not affected at all.

### Progressive gaiting deficiencies in a Purkinje cell-specific model for spinocerebellar ataxia type 1

We next studied a second progressive Purkinje cell degeneration mouse model, the PC-Sca1 mouse, a transgenic mouse line that overexpresses the human *Atxn1* gene with an 82 CAG-repeat under the control of the Purkinje cell-specific *Pcp2* promoter, and that models aspects of the polyQ-disorder spinocerebellar ataxia type 1 (SCA1) (Burright et al., 1995; Clark et al., 1997; White et al., 2021). These PC-Sca1 mice develop progressive gate and balance abnormalities from 5 weeks of age, while eyeblink conditioning, a cerebellar learning paradigm, is already impaired at 4 weeks (Burright et al., 1995; Clark et al., 1997; White et al., 2021; Osório et al., 2023). PC-Sca1 mice also showed progressive reduction in performance in the accelerated rotarod test going from mildly reduced performance at 5 weeks to severely impaired performance at 20 weeks (Clark et al., 1997). The major histopathological hallmark during this time window is the somato-dendritic atrophy of Purkinje cells, starting with subtle loss in dendritic complexity at 3 weeks of age, severe shrinkage of the dendritic tree at 15 weeks of age, and retaining a minimal dendritic tree with small and occasionally heterotopic cell bodies after 25 weeks (Fig. 4A) (Clark et al., 1997; White et al., 2021; Osório et al., 2023). In contrast to PC-Ercc1 KO mice, PC-Sca1 mice maintain largely preserved innervation of the cerebellar nuclei even at 32 weeks of age (Fig. 4B, C). Thus, rather than disappearing, Purkinje cells nerve terminals become smaller and show reduced calbindin staining (Fig. 4B, C). This is consistent with previously published data showing that Purkinje cell loss is limited in PC-Sca1 mice, with 9% loss at 12 weeks and 26-32% loss at 24 weeks of age (Clark et al., 1997; Kaemmerer et al., 2001) (Fig. 4D).

**Figure 4.**
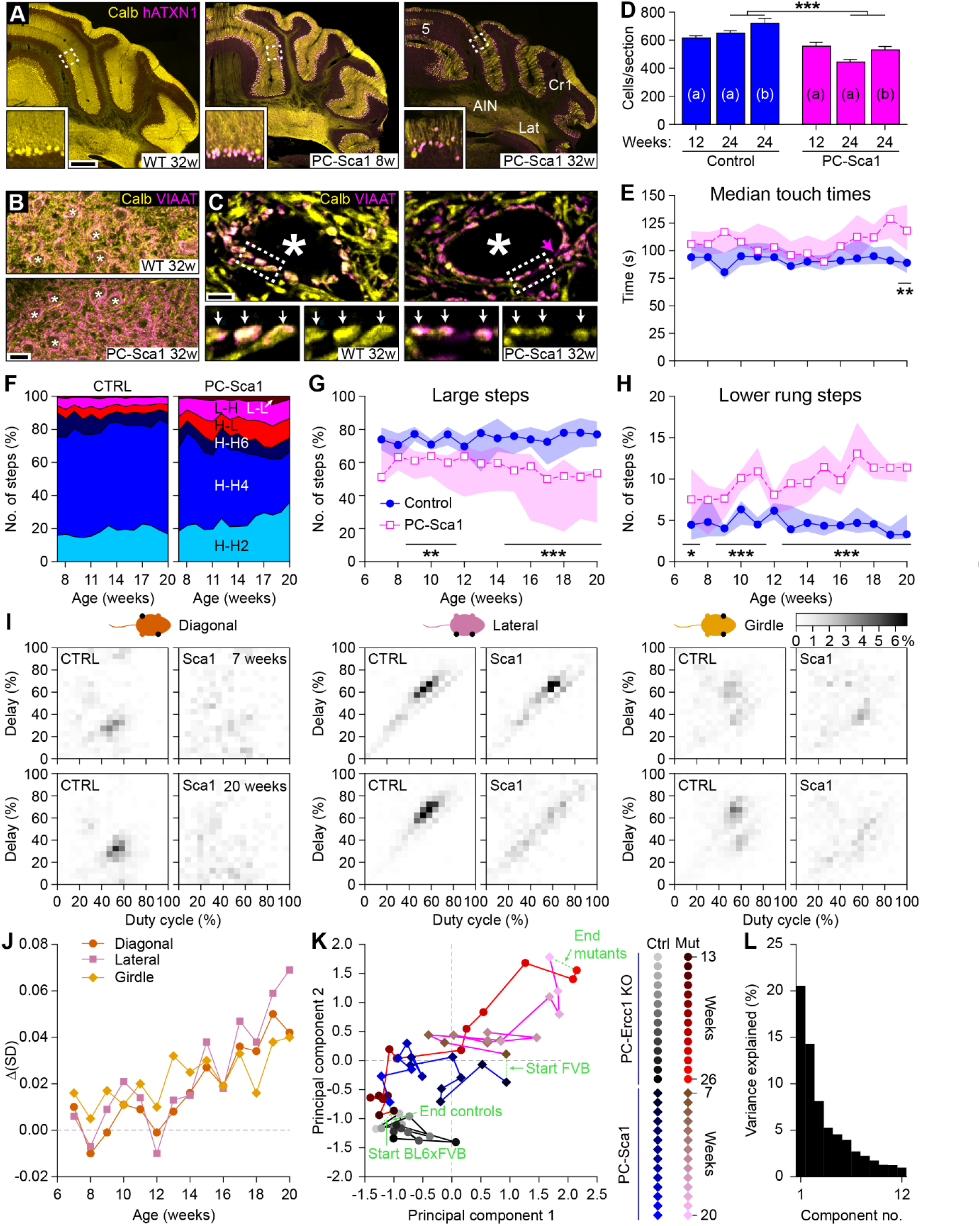
Gait deficiencies in a Purkinje cell-specific mouse model for spinocerebellar ataxia type 1. **A**. Progressive somato-dendritic atrophy and reduced calbindin (Calb) immunostaining of Purkinje cells in the cerebellar cortex. The ATXN1-82Q transgenic protein is selectively labelled with an antibody against human ATXN1 (hATXN1) and is absent in control mice, while strongly expressed in the nuclei of Purkinje cells in transgenic mice. Note, preserved calbindin staining in cerebellar cortex and cerebellar nuclei in PC-Sca1 mice at 8 weeks, and strongly reduced staining in association with Purkinje cell somato-dendritic atrophy at 32 weeks. Overview (**B**) and exemplary cells (**C**) illustrating reduced calbindin immunostaining, but preserved Purkinje cell innervation in the cerebellar nuclei of 32 weeks old PC-Sca1 mice. Note that neurons in the cerebellar nuclei (exemplary cells indicated with an asterisk) are fully surrounded by Purkinje cell axon terminals double labelled for calbindin and VIAAT. However, in 32 weeks old PC-Sca1 mice, Purkinje cell axon terminals are smaller, and show less intense calbindin staining (**C**). Scale bars: 400 µm (**A**), 25 µm (**B**), 5 µm (**C**). **D**. Loss of Purkinje cells PC-Sca1 mice. Graph made from previously published data, counting the number of Purkinje cells per midline sagittal sections, average + SEM. (a) = Clark et al. (1997), (b) = Kaemmerer et al. (2001). Statistics from Clark et al. (1997), comparing WT and PC-Sca1 mice at 12 weeks: *p* = 0.1736; at 24 weeks: *p* = 0.0004, *t* tests. **E**. Touch times. **F**. Distribution of step sizes made by the forelimbs, for abbreviations see Fig. S2G. **F**. Percentage of steps from one high rung to another, with a minimal step size of 4. **G**. Percentage of steps from a higher to a lower rung. Panels D, E, F and H: Median values and interquartile ranges. * *p* < 0.05; *** *p* < 0.001, Mann-Whitney tests with Benjamini-Hochberg correction. **I**. Hildebrand plot representing the combinations of duty cycle (= percentage of step cycle during which the right forelimb touches a rung) and the delay in the step cycle between placement of the right forelimb and the left hindlimb (diagonal), right hindlimb (lateral), and left forelimb (girdle), respectively, at 7 (top) and 20 (bottom) weeks of age. **J**. Changes in the standard deviation of Hildebrand plots. **K**. Principal component analysis of gaiting parameters of PC-Sca1 and PC-Ercc1 KO mice (also see Fig. 3F). The PC-Ercc1 KO mice are at a C57BL/6 x FVB F1 hybrid background and the PC-Sca1 mice at a FVB background, and the latter were younger at the start of the recordings. Note that for both backgrounds, the control and mutant mice of each line start around the same area in the principal component dimensions, and that both mutant lines diverge from their control littermates. At the end of the experiments, the control mice arrive at the same area, suggesting a development and/or training effect for the FVB control mice. Despite their different trajectories and starting points, also both Purkinje cell mutants finally reach similar performance deficits, indicating that, despite changes in the course of the pathology, the end stages are comparable between PC-Ercc1 KO and PC-Sca1 mice **L** Variance explained for each principal component

Like the PC-Ercc1 KO mice, PC-Sca1 mice showed similar trial times in initial sessions as their control littermates, but used more time to travel the ErasmusLadder at more progressed age. This effect was visible at 10 weeks of age, but became statistically significant only in the last session at 20 weeks of age, partially due to the large variation in behavior of the PC-Sca1 mice (Fig. S5B). Similarly, PC-Sca1 mice showed a trend of increased touch times that also only reached statistical significance at 20 weeks of age (Fig. 4E). Notably, the regularity of step times (CV2) was unaltered in PC-Sca1 mice as compared to controls (Fig. S5C). Changes in step size and the incidence of lower rung touches, instead, were significantly different from control mice at earlier ages (Fig. 4F-H), consistent with our findings in PC-Ercc1 KO mice.

As illustrated by the gaiting diagrams of the last session, PC-Sca1 mice, like PC-Ercc1 KO mice, develop more irregular gaiting on the ladder compared to controls (Fig. S5A). Phase analysis of diagonally positioned limbs, indicated that the relative timing of diagonal hindlimb placement versus forelimb was altered in PC-Sca1 mice after 15 weeks (Fig. S5D). Changes in inter-limb coordination in PC-Sca1 mice versus control were substantiated by Hildebrand plots of diagonal, lateral and forelimb pairs with control mice showing stereotyped focal patterns throughout all sessions, and PC-Sca1 mice showing more diffuse distribution of labeled pixels (Fig. 4I,J, Fig. S6). Analysis of variability of pixel intensities of Hildebrand plots indicated changes increased standard deviations indicative of altered inter-limb coordination from about 12-13 weeks for diagonal, lateral as well as forelimb pairs (Fig. 4J). Notably, for all three stances Hildebrand plots from PC-Sca1 mice were identified as deviant from control by a machine learning algorithm from 7 weeks of age (Fig. S6). This suggests that PC-Sca1 mice already at the first session at 7 weeks display subtle changes in inter-limb coordination during gaiting on the ladder (Fig. 5).

**Figure 5.**
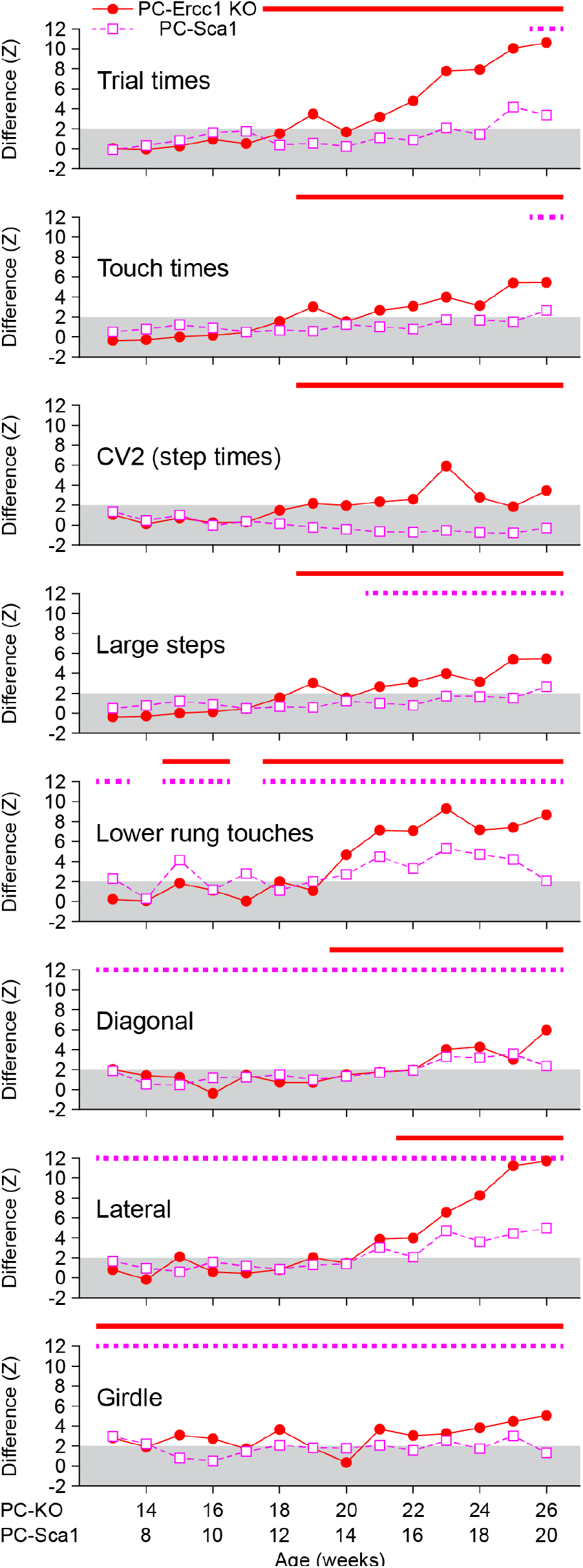
Purkinje cell degeneration can lead to differences in the progression of cerebellar ataxia. Development of the average values of selected gaiting parameters, expressed as the difference between mutant mice and their control littermates. Grey areas indicate 95% confidence interval of control data. While speed, regularity (CV2) and step size are first affected in PC-Ercc1 KO mice, lower rung touches and inter-limb coordination are first affected in PC-Sca1 mice. Notably, the coordination between the right and left forelimb (“girdle”) is the first parameter to be affected in both mouse lines.

Principal component analysis on all gaiting parameters showed a separation of PC-Sca1 mice from control littermates at all ages. During early stages, the PC-Sca1 mice were close to control mice and both primarily moved along the first principal component. From 14-15 weeks onward, PC-Sca1 mice more prominently diverged from the control mice along both the first and second principal components (Fig. S5E,F). Overall principal component analysis indicates that the ErasmusLadder in PC-Sca1 mice, as in PC-Ercc1 KO mice, exposes both mild early changes and progressive late changes in gaiting behavior.

### Subtle differences in progressive gaiting deficiencies in the two Purkinje cell degeneration models

To directly compare the changes in ErasmusLadder gaiting behavior of PC-Ercc1 KO and PC-Sca1 mice we performed principal component analysis. This analysis shows that PC-Ercc1 KO and PC-Sca1 as well as their control littermates move separately in parameter space in initial stages, while developing increasingly convergent paths at later stages, ending closely together (Fig. 4K-L). Significantly, the divergent paths of the mutant lines in the initial stages resemble the paths of the respective control mice. This indicates that differences in initial sessions between PC-Ercc1 KO mice and PC-Sca1, at least in part, result from factors unrelated to the genetic intervention, such as differences in genetic background (C57BL/6J x FVB F1 hybrid versus FVB) and age (13-26 weeks versus 7-20 weeks).

Comparison of Hildebrand plots also indicate that PC-Ercc1 KO and PC-Sca1 mice eventually develop similar locomotor abnormalities on the ErasmusLadder, although they do show differences in the sequence with which these locomotor abnormalities develop. First, subtraction of plots, highlighting how the plots of affected mice differ from their respective control plots, for each combination of limb pairs, reveal similar impact of both mutations studied (Fig. S7A). In addition, graphs plotting the standard deviation of pixel intensities of the consecutive sessions, again for each combination of limbs, show overlapping trajectories for PC-Ercc1 KO versus PC-Sca1 mice when aligned for session number (Fig. S7B). Hildebrand plots for diagonal and lateral stance showed relatively unaltered variations in the initial sessions, while showing increased standard deviations starting from about halfway the testing period (20 and 14 weeks for Ercc1-PC KO versus PC-Sca1 mice, respectively). Instead, girdle, thus the coordination between the two forelimbs, increased standard deviation that persisted throughout the testing period, and that was already visible at the first sessions of both mutants. With unsupervised machine learning, however, early changes in diagonal and lateral stance could already be observed early on during the disease process in PC-Sca1 mice, and only much later in PC-Ercc1 KO mice.

Comparison of gaiting parameters from PC-Ercc1 KO versus PC-Sca1 mice expressed as Z-scores from their respective controls, point to some differences between the two lines (Fig. 5). Most notably, in PC-Sca1 mice CV2 of step times were unaltered compared to controls, while strongly increased compared to control in late phase PC-Ercc1 KO mice (Fig. 5, Fig.S2E, Fig. S5A). Another difference is that while PC-Ercc1 KO mice show relatively robust progressive changes in multiple parameters from about 18 weeks of age, in PC-Sca1 changes are less robust and develop more gradually (Fig. 5). Importantly, in both lines the earliest changes were increased frequency of lower rung touches and reduced coordination of forelimbs (Fig. 2E, 4H, 5). In summary, PC-Ercc1 KO and PC-Sca1 mice eventually develop similar changes in locomotor parameters, but follow different sequences in reaching that stage.

## Discussion

Gait ataxia is a dominant feature of cerebellar disorders, and detailed gait analysis can be used in the clinic to follow the disease progression from prodromal to late stages (Jacobi et al., 2013; Rochester et al., 2014; Velázquez-Pérez et al., 2014a; Velázquez-Pérez et al., 2014b; Maas et al., 2015; Ilg et al., 2016; Velázquez-Pérez et al., 2017; Kim et al., 2021; Shah et al., 2021; Velázquez-Pérez et al., 2021; Thierfelder et al., 2022). The ability to follow changes in the gaiting pattern of mouse models for cerebellar ataxia, starting from the very first signs, is vital to obtain further insight into pathological mechanisms, and for the development and validation of intervention (Cendelin et al., 2022). Following up on our findings that gait analysis on an automated horizontal ladder, the ErasmusLadder, provides a spectrum of changed locomotor parameters that allowed to differentiate between cell-specific cerebellar mouse models with deficits of different severity (Vinueza Veloz et al., 2015), here we present a longitudinal study using the ErasmusLadder to monitor the development of locomotor abnormalities in two mouse lines with different types of adult-onset progressive Purkinje cell degeneration. The PC-Ercc1 KO mice are mainly characterized by a stochastic loss of Purkinje cells. Following a moderate loss of Purkinje cells, these mice rapidly develop ataxic gait, including slower and shorter steps, with more lower rung touches and less coordination between the four limbs. In contrast, the PC-Sca1 mice initially mainly show atrophy of Purkinje cells, with limited Purkinje cell loss and intact innervation of the cerebellar nuclei (Ingram et al., 2016; White et al., 2021). From early on, they show a lack of inter-limb coordination and increased usage of the lower rungs, followed by smaller and slower steps. Remarkably, the regularity of the step timing is maintained in the PC-Sca1 mice. Thus, the two mouse lines displaying different forms of Purkinje cell degeneration also show qualitative and quantitative differences in progression of locomotor changes on the Erasmusladder. In other words, cerebellar ataxia is not necessarily a gradual and sequential loss of specific locomotor capabilities, but follows a patterned deterioration that depends on the pathological trigger.

### Comparison with human diseases

Although dysfunction, disconnection and eventual loss of Purkinje cells are central to many degenerative forms of cerebellar ataxia, extracerebellar damage can be substantial in patients and knock in mouse models (Hekman and Gomez, 2015; Huang and Verbeek, 2019; Klockgether et al., 2019; Sen et al., 2019; Coarelli et al., 2023). Thus, it is generally not straightforward to link Purkinje cell pathology to the development of ataxic gait in patients. To isolate the impact of adult-onset Purkinje cell degeneration on the development of ataxic gait, we have studied here two mouse models with mutations restricted to Purkinje cells. Despite differences between the bipedal gait of humans and the quadrupedal gait of mice, we observed slower, smaller and uncoordinated steps in our mouse models, just as in patients. Loss of Purkinje cell function can, therefore, by itself explain the development of ataxic gait. Clear signs of tremor and dystonia, which both can occur in patients with cerebellar damage (Diener et al., 1984; Deuschl et al., 2000; Neychev et al., 2008; Shakkottai et al., 2017; Nikolov et al., 2019; Dankova et al., 2020; Manto et al., 2020; Pan et al., 2020; Giannì et al., 2022; Klomp et al., 2022; Brown et al., 2023), were absent in our mice. Although we did not specifically test for tremor or dystonia, so that subtle signs may have been overlooked, this may imply that tremor and dystonia rely less on the contribution of malfunctioning Purkinje cells than does ataxic gait.

Late-onset cerebellar ataxia is often considered to be a degenerative disease affecting older patients. However, brain scans have documented subtle changes in the shape of the human cerebellum at the prodromal stage of cerebellar ataxia (Jacobi et al., 2013; Velázquez-Pérez et al., 2014b; Maas et al., 2015; Nigri et al., 2020; Kim et al., 2021). Accordingly, many symptoms, including gait and stance deficits, can already be observed before the clinical diagnosis of cerebellar ataxia is established (Jacobi et al., 2013; Velázquez-Pérez et al., 2014a; Velázquez-Pérez et al., 2014b; Maas et al., 2015; Velázquez-Pérez et al., 2017; Kim et al., 2021; Velázquez-Pérez et al., 2021). Similarly, animal experiments have shown that eyeblink conditioning, a form of procedural learning critically depending on the cerebellum, can already be impaired in the pre-ataxia stage (Osório et al., 2023). Together with our current data, it becomes clear that many forms of cerebellar ataxia can already become manifest before overt clinical signs occur. Consequently, future therapeutic intervention might be optimally targeted at prodromal or early stages of the disease. For this reason, sensitive tests for ataxic gait in mouse models, as described in this study, will be valuable.

### Cerebellar ataxia in mouse models

Our data show that the ErasmusLadder is a highly valuable tool for longitudinal studies of locomotor deficiencies. The automated nature of the ErasmusLadder, in combination with the many repetitions and a relatively challenging motor task, leads to well-quantifiable data on mouse locomotion that allows identification of relatively subtle changes in behavior. As the description of the gaiting pattern relies on the interpretation of sensor touches, post-hoc data processing is required to interpret limb placement. This interpretation gets more difficult when the gaiting pattern becomes very irregular in progressed ataxia, and mice may touch the rungs also with their snout or belly. We show here, in line with earlier descriptions of at least moderate forms of ataxia (Van Der Giessen et al., 2008; Vinueza Veloz et al., 2015), that also in these cases the ErasmusLadder can still describe gaiting deficits, but the level of detail that can be retrieved decreases in severely affected individuals. We argue, therefore, that the ErasmusLadder is particularly useful at prodromal and early disease stages.

Despite the use of different background strains, both mouse models reach similar levels of gait ataxia. The comparison between the two wild type groups emphasizes the use of the correct controls. In particular, FVB mice are albino and thus have probably poor eyesight (van Wyk et al., 2015). From the start on, they have longer touch times, more lower rung touches and a higher CV2 of the step times than C57BL/6 mice.

### Different approaches to measure locomotion deficits

The new analysis methods introduced in this study allow us to retrieve the placement of all four limbs, a feature that was not readily available previously for the ErasmusLadder (Vinueza Veloz et al., 2015). This enabled us to construct Hildebrand plots, pioneered for the study of horse gaiting and later applied to several other species (Hildebrand, 1965; Cartmill et al., 2020; Struble and Gibb, 2022). Possibly related to the relatively short gangway of the ErasmusLadder, Hildebrand plots from individual mice turned out to be too noisy for further analysis, so that we had to group all steps per session and genotype. Nevertheless, this analysis turned out to be powerful and insightful: when combined with unsupervised machine learning, subtle changes in inter-limb coordination could be revealed. It was interesting to note that the coordination between right and left forelimbs was affected before that between the fore- and hindlimbs at the same side in the PC-Ercc1 KO mice, suggesting a relatively strong contribution of the cerebellum to bilateral coordination (see also (2022)).

Other tests for gait ataxia exist. Of these, the accelerating rotarod is probably the most widely used (Dunham and Miya, 1957; Jones and Roberts, 1968; Lalonde et al., 1995; Tsai et al., 2012). Like the ErasmusLadder, the accelerating rotarod is a physically challenging task for mice, but it yields only one read out – latency to fall – per trial, instead of hundreds of timed rung touches with the ErasmusLadder. As a consequence, the accelerating rotarod is not very sensitive to relatively subtle cerebellar deficits (Stroobants et al., 2013). In the present study, we noticed that the ErasmusLadder could identify gait ataxia in PC-Ercc1 KO mice more than two months before the accelerating rotarod. In contrast, for the PC-Sca1 mice, the rotarod was approximately equally sensitive to early changes in behavior as the ErasmusLadder (Clark et al., 1997).

The beam walk test can be an alternative or complementary test. It reportedly has a higher sensitivity for cerebellar deficits than the rotarod (Stanley et al., 2005). The PC-Sca1 mice used in the current study displayed an overt phenotype on the beam walk at 7 weeks of age, thus comparable with the ErasmusLadder (White et al., 2021). It can be assumed that the vestibular system is very important for successful trials on the beam walk, making the interpretation of beam walk results in terms of locomotor control per se more difficult.

In addition, footprint analysis can be used, for instance in conjunction with the CatWalk (Timotius et al., 2019; Heinzel et al., 2020; Mészáros et al., 2021). This test yields a detailed impressions of the exact limb placement and force used during each step, but it relies on spontaneous locomotion, resulting in less and more variable trials than the ErasmusLadder. Furthermore, with the recent advances in video analysis, high-speed videography is bound to become an important addition to gait analysis (Machado et al., 2015; Mathis et al., 2018; Machado et al., 2020; Liu et al., 2021). Yet, the ErasmusLadder test is designed to have a large number of repetitions of a relatively challenging locomotor task together with a strong motivation (tail and head winds) to perform optimally. It is still an unfulfilled challenge to combine such a task with video analysis.

Thus, while we argue here that the ErasmusLadder provides a sensitive test for cerebellar ataxia, alternative tests for the quantification of cerebellar ataxia in mice are available. In general, tasks that are not physically challenging and/or have only a few repetitions per session, are likely to be less sensitive, so that the absence of a phenotype on such a task should be considered with caution.

### Conclusions

Insight into the relation between Purkinje cell dysfunction and patterned loss of locomotor abilities allows us to study differential contributions of Purkinje cells to specific aspects of motor control. The ErasmusLadder provides thereby a sensitive paradigm for testing subtle changes in gait patterns over time, and can, therefore, used for future intervention studies aimed at prodromal and early disease stages of cerebellar ataxia and other motor diseases.

## Acknowledgements

The authors wish to thank Stephanie Dijkhuizen, Laura Post, Renate Brandt, Alexander Cupido and Elise Haasdijk for technical assistance, Lorenzo Bina and Lucas Veeger for contributing to the pilot phase of this study, Reinko Roelofs for conceptual discussions, and Jan Hoeijmakers and Ingrid van der Pluijm for providing the PC-Ercc1 KO mice.

