## Supplemental data for "Different Purkinje cell pathologies cause specific patterns of progressive ataxia in mice"

### Supplementary figures

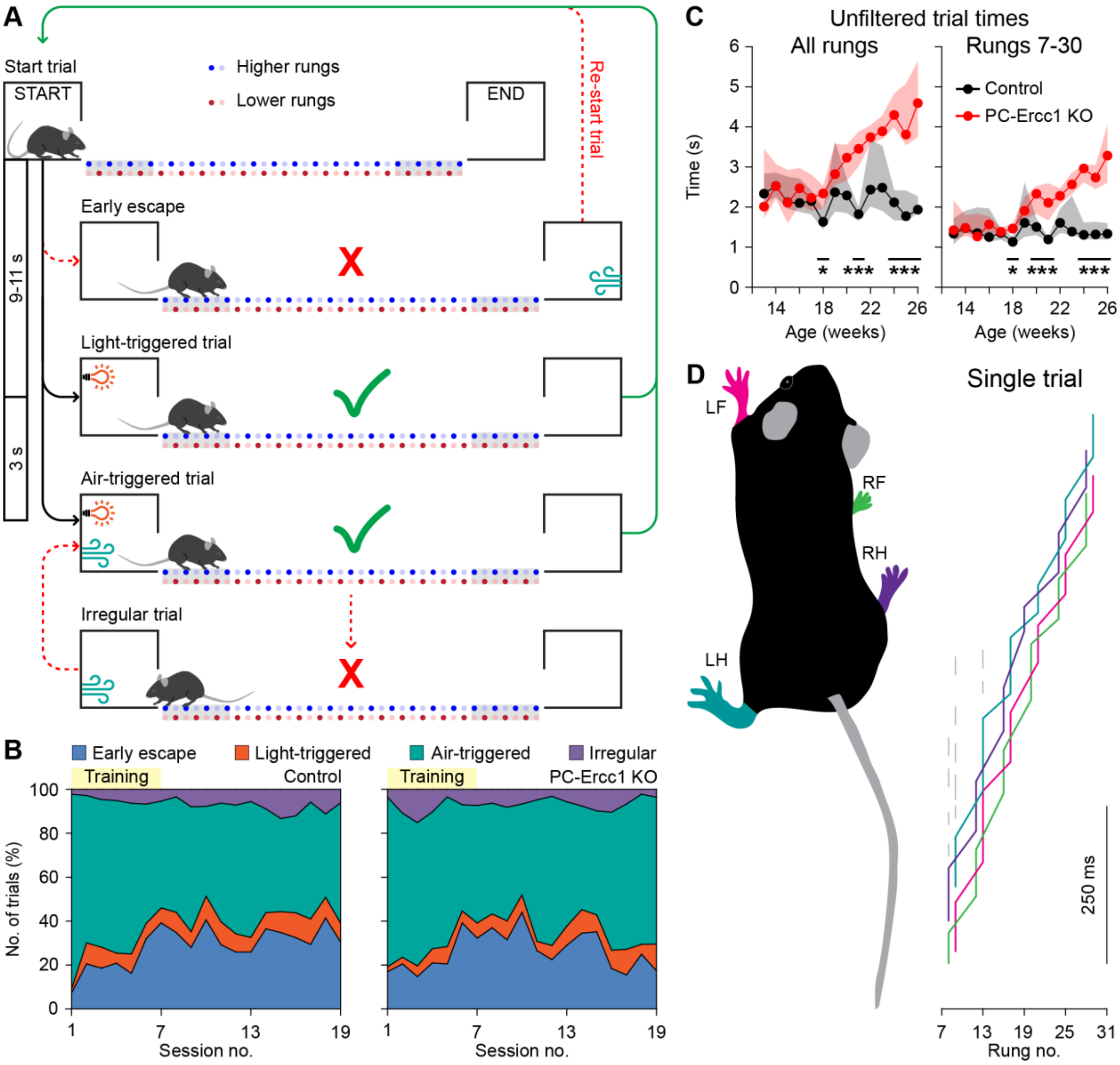

**Figure S1. Setup and analysis of locomotion on the ErasmusLadder**

**A.** The start of a trial was announced by lighting an LED in the start box. Early escapes, i.e., departures from the start box before the LED was turned on, were discouraged by a strong head-wind. Early escapes induced the abortion of a trial, and that trial was then repeated until the mice left after lighting the LED. Data form early escapes were not included in the analysis of gaiting. Three seconds after the LED was turned on, a strong tail-wind encouraged the mouse to walk efficiently to the end box. Returns to the start box occasionally occurred, and the steps made during these unfinished crossings were excluded from the analysis of gaiting. 9-11 seconds after arriving in the end box, the next trial started, during which the mice crossed the ErasmusLadder in the opposite direction. **B.** The majority of trials started as air-triggered trials (see A), in both PC-Ercc1 KO and control mice. **C.** Unfiltered trial times (see Methods) from start to end (*left*) and on middle part (*right*). Medians and inter-quartile ranges. * $p < 0.05$; *** $p < 0.001$, Mann-Whitney tests with Benjamini-Hochberg correction. **D.** Using post-hoc analysis, we determined the most likely placement of the four limbs on the ErasmusLadder. Each limb is color-coded (*left*). Representative single trial of a control mouse (*right*). Vertical line segments represent the stance phase, the diagonal parts the swing phase, and the colors refer to the four limbs. The grey lines indicate registered sensor activity that was not attributed to any of the limbs.

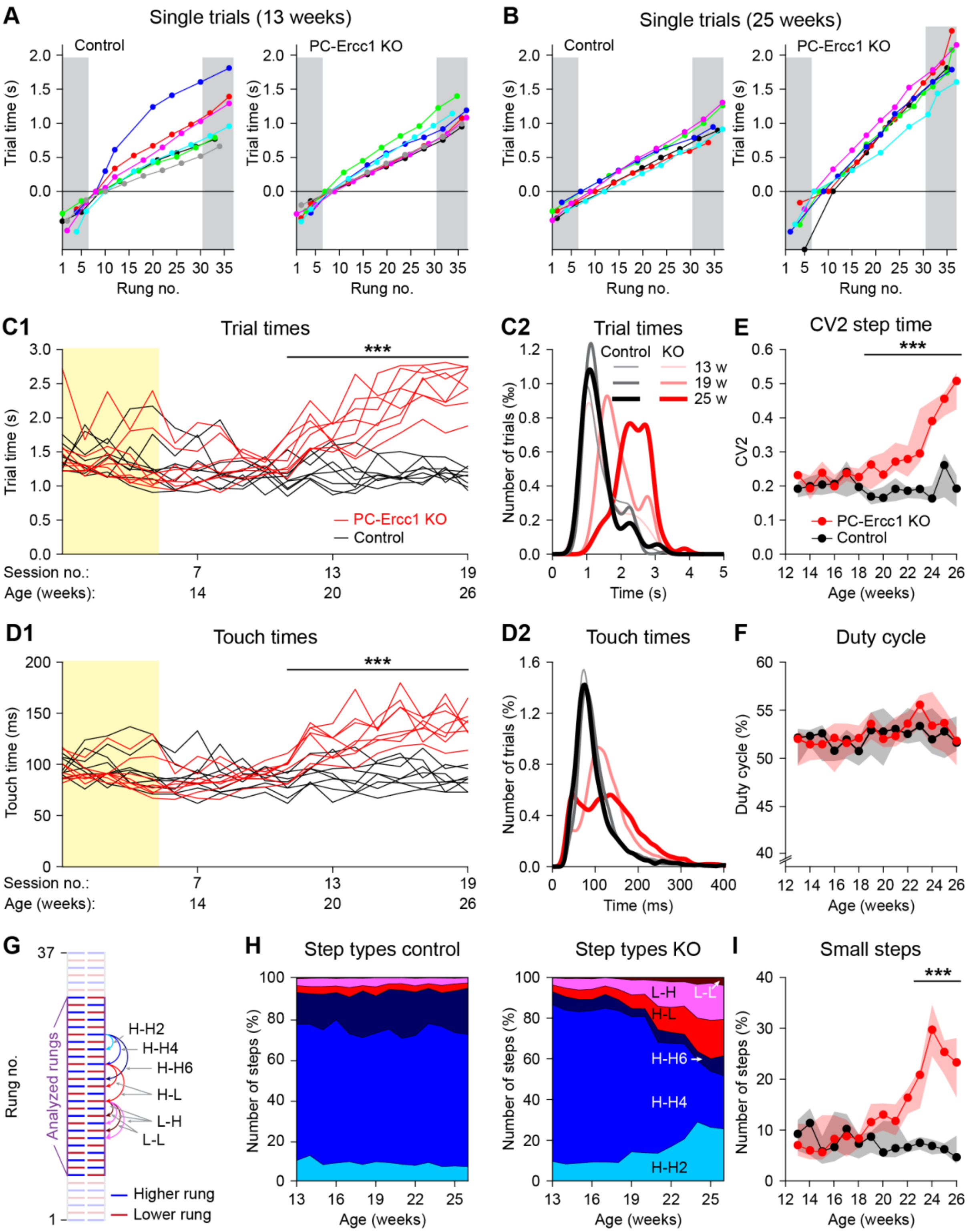

**Figure S2. Slower, shorter and less accurate steps in PC-Ercc1 KO mice**

**A.** Of each mouse, we plotted the trajectories of the right forelimb during the first regular trial at 13 weeks of age. The grey strips on the side indicate the first and last rungs that were excluded from subsequent gaiting analysis. Hence, the first touch on the first rung after rung no. 6 was considered the moment of trial start. **B.** As A, but at 25 weeks of age. Note the slower and less regular gaiting in the PC-Ercc1 KO mice. **C1.** Trial times for each individual mouse, including the training period (sessions 1-5 at 12 weeks of age). **C2.** Histograms of the trial times at 13, 19 and 25 weeks of age. **C1.** Touch times of individual mice. **C2.** Histograms of the touch times at 13, 19 and 25 weeks of age. **D.** Local variation (CV2) of the step times. **E.** Percentage of the step cycle during which the right forelimb was touching a rung. **G.** Schematic drawing of the rungs on the ErasmusLadder, showing the alternating pattern of higher and lower rungs. The rungs 1-6 and 31-37 were not analyzed when considering the gaiting pattern. The arrows on the side indicate the possible step types, with the letter indicating higher (H) and lower (L) rungs and the number the step size. **H.** Surface plot indicating the distribution of step types in control and PC-Ercc1 KO mice over time. It is obvious that the number of large steps decreased (see also Fig. 2D) and that of lower rung touches increased over time in PC-Ercc1 KO mice (see also Fig. 2E). **I.** Increase in number of small steps (from one higher rung to the next, H-H2). Panels **E**, **F** and **I** illustrate medians and interquartile ranges. * $p < 0.05$, *** $p < 0.001$, Mann-Whitney tests with Benjamini-Hochberg correction.

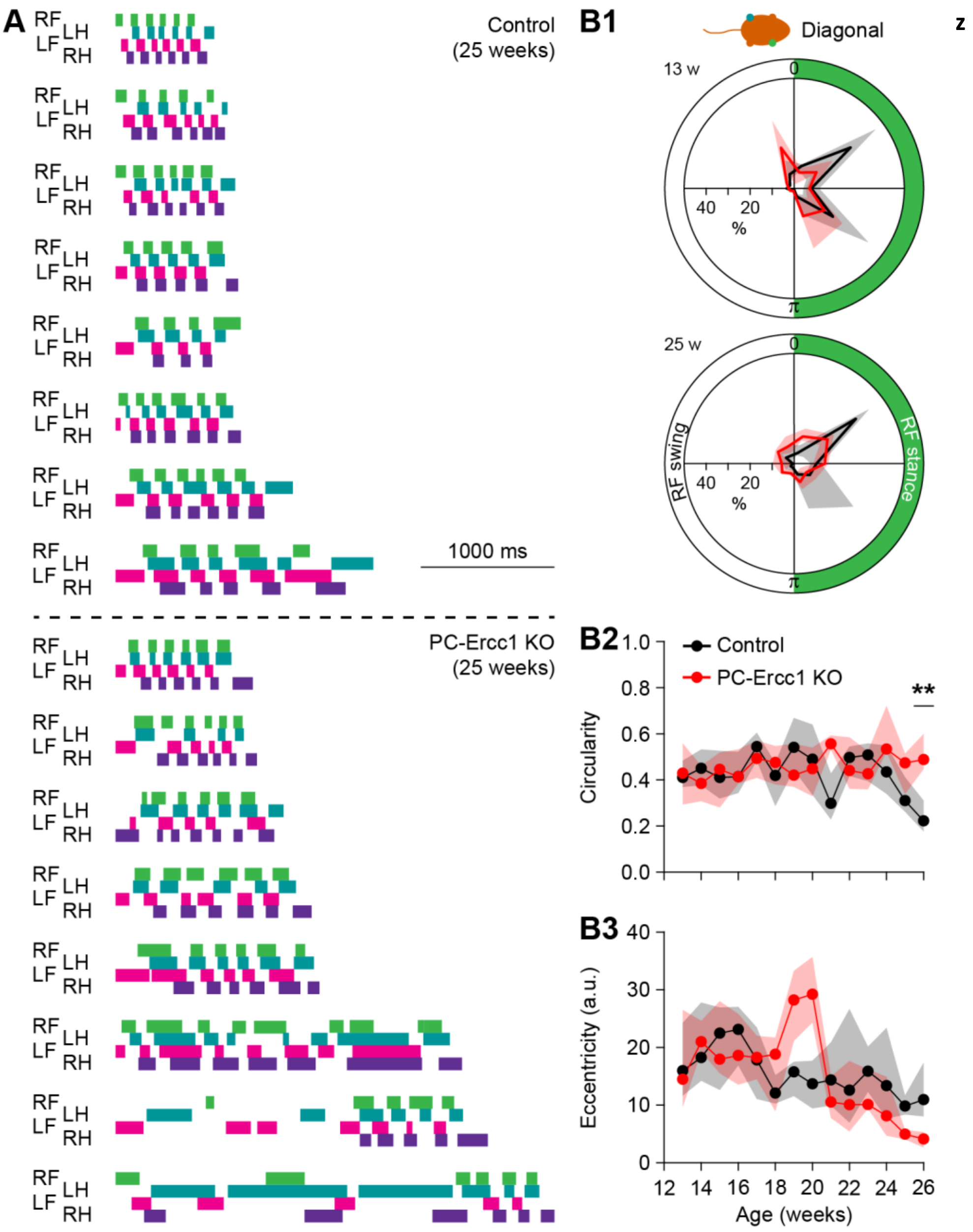

**Figure S3. PC-Ercc1 KO mice show decreased intra-limb coordination over time**

**A.** Representative gait plots illustrating stance (filled bars) and swing (open spaces in between) phases of control and PC-Ercc1 KO mice at 25 weeks. Plots are based on the first regular trial of each mouse. For this illustration, the whole trials were plotted, thus including the first and last rung that were typically excluded from quantitative analysis. **B1.** Phase analysis of the left hindlimb in comparison to the step cycle of the right forelimb. The step cycle is divided into two equal parts, determined by the placement (0) and lift (π). The polar plots indicate the phases of placement of the left hindlimb. Note, two peaks of hindlimb placement in the polar plot of control mice at both 13 and 25 weeks (black line with grey shading), and three peaks in the polar plot of PC-Ercc1 KO mice at 13 weeks (red line and shading). At 25 weeks, the PC-Ercc1 KO mice lost most of their pattern in this diagonal stance indicated by the absence of peaks. However, circularity (**B2**) and eccentricity (**B3**) of the polar plots in **B1,** were not show consistent changes in time in PC-Ercc1 KO versus control mice. ** $p < 0.01$, Mann-Whitney tests with Benjamini-Hochberg correction.

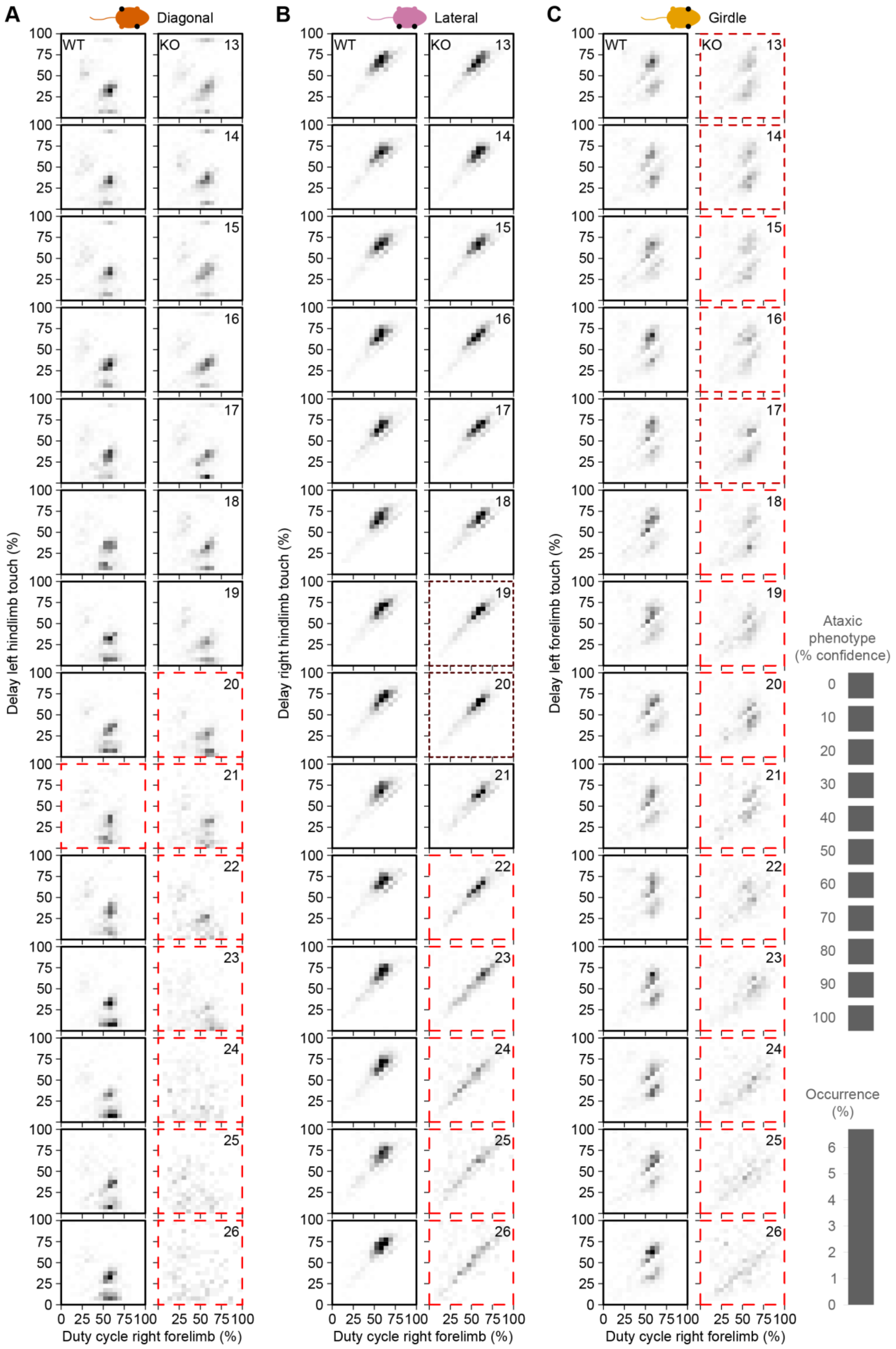
A
Diagonal
B
Lateral
C
Girdle
WT
KO
Delay left hindlimb touch (%)
Delay right hindlimb touch (%)
Delay left forelimb touch (%)
Duty cycle right forelimb (%)
Ataxic phenotype (% confidence)
Occurrence (%)

**Figure S4. PC-Ercc1 KO mice show impaired inter-limb coordination.**

**A.** Hildebrand plots representing the combinations of duty cycle (= percentage of step cycle during which the right forelimb touches a rung) and the delay in the step cycle between placement of the right forelimb and the left hindlimb. All registered steps from all mice were grouped per session and genotype. The same for the comparison between the duty cycle from the right frond limb and the right hindlimb (**B**) or the left forelimb (**C**). The thick colored borders indicate the sessions that were marked as deviant by unsupervised machine learning (see Methods).

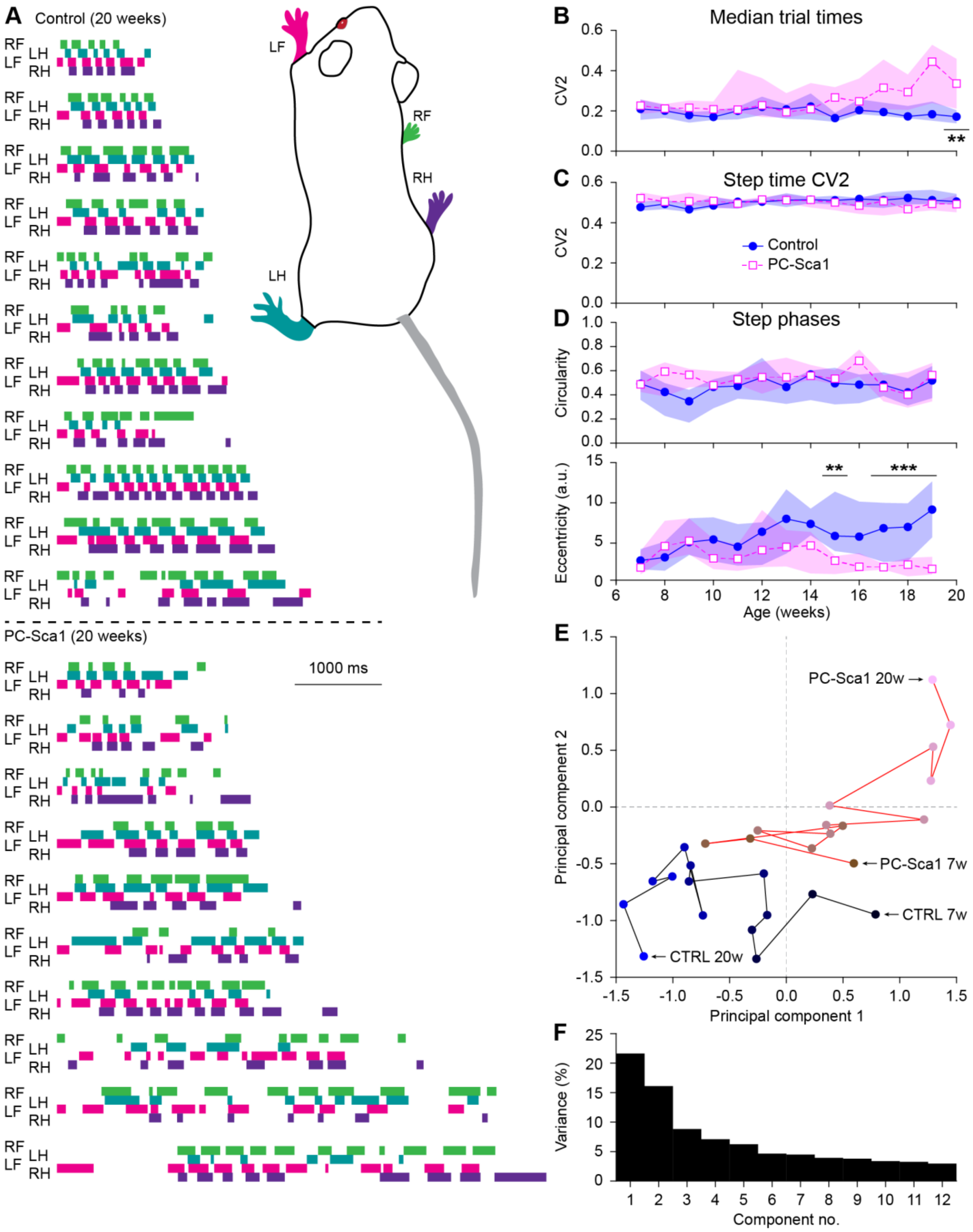

**Figure S5. PC-Sca1 mice show decreased intra-limb coordination over time**

**A.** For each mouse, the first regular trial at 20 weeks of age was plotted. The filled bars indicate rung touch (stance phase) and the open spaces in between the absence of sensor touch (swing phase). For this illustration, the whole trials were plotted, thus including the first and last rung that were typically excluded from quantitative analysis. **B.** Trial times (rungs 7-30). **C.** CV2 of the step times was not affected in PC-Sca1 mice. **D.** Phase analysis of the left hindlimb in comparison to the right forelimb (see also Fig. S3D). Medians with interquartile range. **E.** Principal component analysis reveals that both groups of mice initially show comparable behavior, that diverges later on. **F.** Variance explained for each principal component. *** $p < 0.001$, Mann-Whitney tests with Benjamini-Hochberg correction.

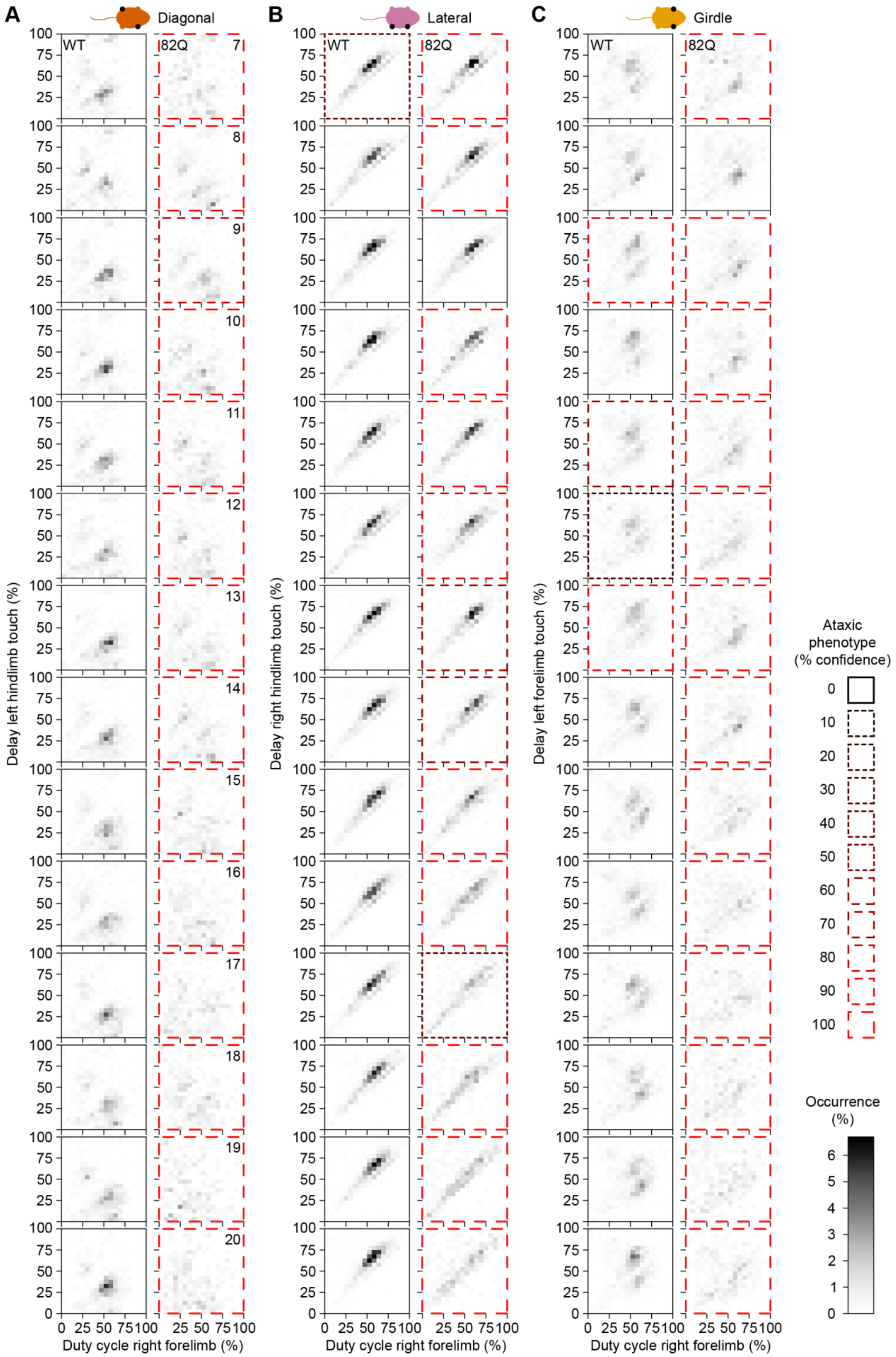

A
Diagonal
B
Lateral
C
Girdle
WT
82Q
Delay left hindlimb touch (%)
Delay right hindlimb touch (%)
Delay left forelimb touch (%)
Duty cycle right forelimb (%)
Ataxic phenotype (% confidence)
0
10
20
30
40
50
60
70
80
90
100
Occurrence (%)

**Figure S6. PC-Sca1 mice show impaired inter-limb coordination.**
**A.** Hildebrand plots representing the combinations of duty cycle (= percentage of step cycle during which the right forelimb touches a rung) and the delay in the step cycle between placement of the right forelimb and the left hindlimb. All registered steps from all mice were grouped per session and genotype. The same for the comparison between the duty cycle from the right frond limb and the right hindlimb (**B**) or the left forelimb (**C**). The thick colored borders indicate the sessions that were marked as deviant by unsupervised machine learning (see Methods). 82Q refers to the PC-Sca1 mice.

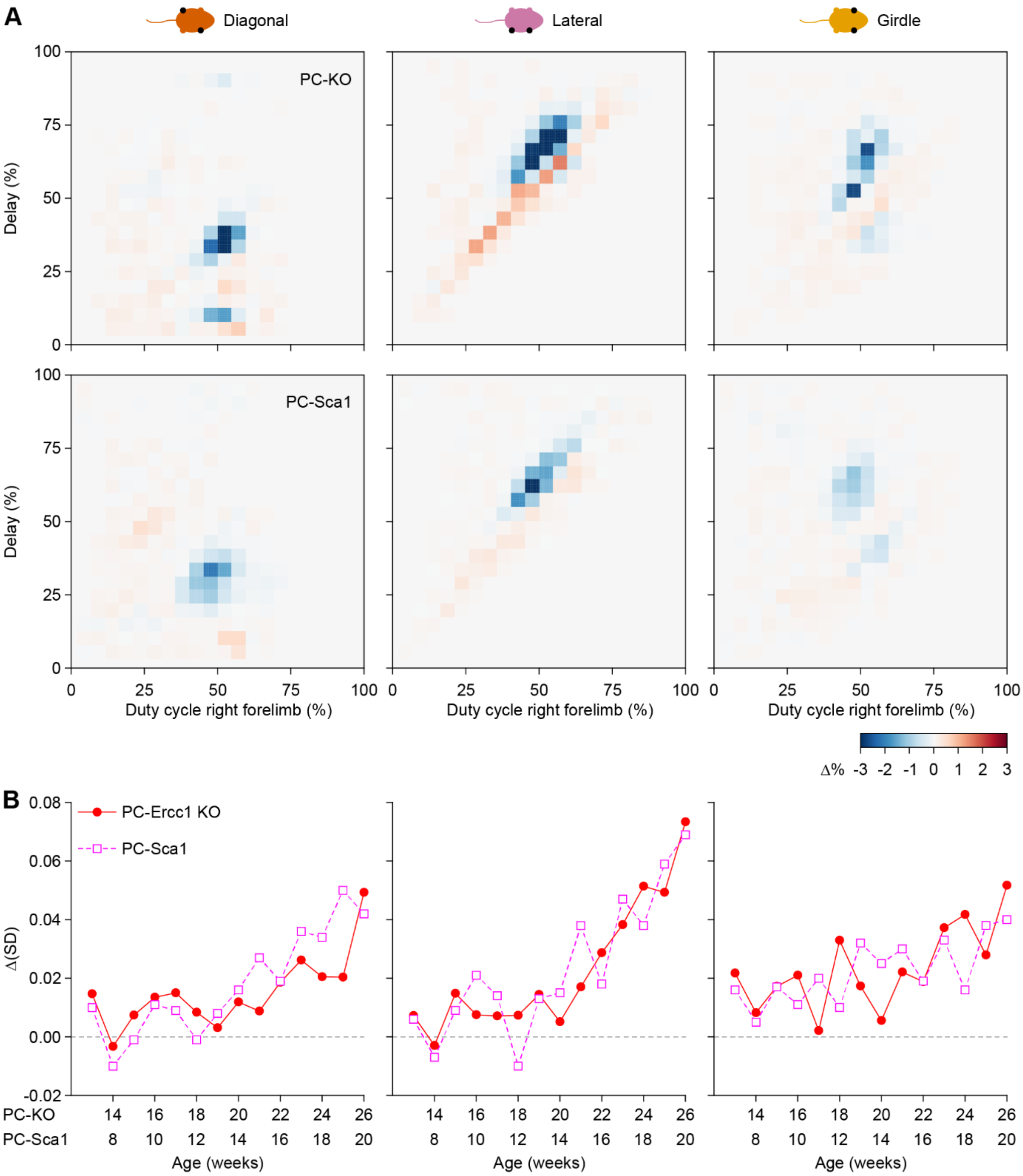

**Figure S7. Intralimb coordination is affected by Purkinje cell degeneration**

**A.** The intralimb coordination analysis of Fig. S4 and Fig. S6 were subjected to unsupervised machine learning, that defined, for each combination of limbs (see icons above the graphs) and each mouse strain, the difference in density of occurrences of gaiting patterns between the clusters separated as control and deviant behavior. Blue indicates more WT-like behavior, red more ataxic behavior. The upper row is copied from Fig. 3E to facilitate comparison between the two strains. Despite their different background strains, age of onset, and course of disease, the differences in patterns are strikingly similar between the two mouse lines. **B.** Standard deviations of the pixel intensities of the plots of Fig. S4 and Fig. S6, showing similar trajectories in losing intralimb coordination. Note that only for diagonal placement, the PC-Sca1 showed an earlier effect than the PC-Ercc1 KO mice.

**Table S1 – Summary of previously published studies characterizing mouse locomotion during unperturbed trials on the ErasmusLadder**

| Mutation [background] | Model for | Parameter | Result | Reference |
|---|---|---|---|---|
| *Gjd2* KO [C57BL/6] | Electrotonic coupling | No. of lower rung touches | No effect | Van Der Giessen et al., 2008 |
| | | Step time | No effect | |
| Lurcher [C57BL/6] | Cerebellar degeneration | No. of lower rung touches | Increased | Van Der Giessen et al., 2008 |
| | | Step time | No effect | |
| | Cerebellar degeneration | Step time | Increased | Renier et al., 2010 |
| | | Successful trials[1] | No effect | |
| *Ptf1a-Cre;Robo3*$^{lox/lox}$ [C57BL/6] | HGPPS | Step time | Increased | Renier et al., 2010 |
| | | Successful trials[1] | Decreased | |
| PICK1 [C57BL/6] | LTD deficiency | No. of lower rung touches | No effect | Schonewille et al., 2011 |
| | | Step time | No effect | |
| GluR2Δ7 [C57BL/6] | LTD deficiency | No. of lower rung touches | No effect | Schonewille et al., 2011 |
| | | Step time | No effect | |
| GluR2K882A [C57BL/6] | LTD deficiency | No. of lower rung touches | No effect | Schonewille et al., 2011 |
| | | Step time | No effect | |
| *Nf1*$^{+/-}$ [C57BL/6] | Neuro-fibromatosis type 1 | No. of lower rung touches | No effect | van der Vaart et al., 2011 |
| | | Step time | No effect | |
| *Nlgn3*$^{KO}$ [C57BL/6] | Autism spectrum disorder | No. of lower rung touches | No effect | Baudouin et al., 2012 |
| | | Step time | Increased | |
| *Nlgn3*$^{PC}$ [2] [C57BL/6] | Autism spectrum disorder | No. of lower rung touches | Increased | Baudouin et al., 2012 |
| | | Step time | Increased | |
| GLAST$^{CreERT2/+}$ × Gria1$^{fl/fl}$ × Gria4$^{fl/fl}$ [C57BL/6] | Bergman glial cell function | No. of lower rung touches | No effect | Saab et al., 2012 |
| *Fmr1* KO [FVB/Ant] | Fragile X syndrome | Step time | No effect | Vinueza Veloz et al., 2012 |
| *Gabra6*-Cre;*Cacna1a* KO [C57BL/6] | Granule cell function | No. of lower rung touches | No effect | Galliano et al., 2013 |
| *Gabra6*-Cre;*Cacna1a* KO [C57BL/6J] | Granule cell function | No. of lower rung touches | No effect | Vinueza Veloz et al., 2015 |
| | | Step size | No effect | |
| | | Step time | No effect | |
| | | Step time CV2 | No effect | |
| *Npc1*$^{-/-}$ [Balb/c] | Niemann-Pick type C | Step time | Increased | Marques et al., 2015 |

| | | | | |
|---|---|---|---|---|
| Pcd [C57BL/6J] | Cerebellar degeneration | No. of lower rung touches | Increased | Vinueza Veloz et al., 2015 |
| | | Step size | Decreased | |
| | | Step time | Increased | |
| | | Step time CV2 | Increased | |
| *Pcp2-Ppp3r1 (L7-PP2B)* [C57BL/6J] | LTP deficiency | No. of lower rung touches | No effect | Vinueza Veloz et al., 2015 |
| | | Step size | Decreased | |
| | | Step time | Increased | |
| | | Step time CV2 | No effect | |
| *Pcp2-Gabrg2* KO *(L7-Δγ2)* [C57BL/6J] | Molecular layer interneuron function | No. of lower rung touches | No effect | Vinueza Veloz et al., 2015 |
| | | Step size | No effect | |
| | | Step time | No effect | |
| | | Step time CV2 | No effect | |
| *Shank2* KO [C57BL/6N] | Autism spectrum disorder | No. of lower rung touches | Increased | Ha et al., 2016 |
| | | No. of backsteps and jumps | Increased | |
| *Pcp2-Shank2* KO [C57BL/6N] | Autism spectrum disorder | No. of lower rung touches | Increased | Ha et al., 2016 |
| | | No. of backsteps and jumps | Increased | |
| *Pcp2-Shank2* KO [C57BL/6] | Autism spectrum disorder | Step time | No effect | Peter et al., 2016 |
| *Pcp2-Slc26a11* KO [C57BL/6] | Chloride homeostasis | Step size | Decreased | Rahmati et al., 2016 |
| *Pcp2-Foxp2* KO[C57BL/6J] | "Speech disorders" | No. of lower rung touches | Increased | French et al., 2018 |
| Sox14-positive cell ablation [C57BL/6] | Nucleo-olivary projections | No. of lower rung touches | Increased | Prekop et al., 2018 |
| | | Step time | No effect | |
| | | No. of jumps | Decreased | |
| Hypoxic treatment [C57BL/6] | Hypoxia | No. of lower rung touches | Increased | Sathyanesan et al., 2018 |
| | | No. of backsteps | Increased | |
| | | Step time | No effect | |
| Tg-rmd [C57BL/6J] | MDCMC | No. of lower rung touches | No effect | Sayed-Zahid et al., 2019 |
| | | No. of backsteps | No effect | |
| *Pcp2-TrpC3* KO [C57BL/6J] | Purkinje cell heterogeneity | No. of steps | No effect | Wu et al., 2019 |
| SCD mice (sickle cell) (Townes mice) [C57BL/6] | Sickle cell disease | No. of lower rung touches | Increased | Almeida et al., 2020 |
| | | Step size | Complex phenotype | |
| | | No. of backsteps | No effect | |
| | | Step time | No effect | |
| *Pcp2-SK2* KO [C57BL/6J] | Purkinje cell excitability | No. of lower rung touches | No effect | Grasselli et al., 2020 |
| | | Step size | Decreased | |

| | | | | |
|---|---|---|---|---|
| *Fmr1[103Q]* [C57BL/6J] | FXTAS | No. of lower rung touches | No effect | Haify et al., 2020 |
| | | Step size | No effect | |
| | | Step time | No effect | |
| Traumatic brain injury [ICR] | Traumatic brain injury | No. of lower rung touches | Increased | Namdar et al., 2020 |
| | | Trial duration | No effect | |
| *Pcp2-Shisa6* KO [C57BL/6] | Autism spectrum disorder | No. of lower rung touches | Increased | Peter et al., 2020 |
| *Sphk1*$^{-/-}$ [C57BL/6J] | Involved in SCA1 | Step size | No effect | Blot et al., 2021 |
| *Pcp2-tau-eGFP* [C57BL/6J] | Axonal swellings | No. of short steps | [3] | Lang-Ouellette et al., 2021 |
| TRE-36×$G_4C_2$-GFP/hnRNP-rtTA DT [C57BL/6JRj] | C9ORF72 | No. of lower rung touches | Increased | Riemslagh et al., 2021 |
| ckrc14$^{CYP19a}$ [C57BL/6] | Placental dysfunction | No. of lower rung touches | No effect | Vacher et al., 2021 |
| | | Step size | Decreased | |
| ATXN1[82Q] [FVB/N] | SCA1 | No. of lower rung touches | No effect | White et al., 2021 |
| | | Step size | Decreased | |
| *Pax5*$_{+/-}$; *Pax5*$^{R31Q/-}$ [C57BL/6] | Autism spectrum disorder | No. of lower rung touches | Increased | Kaiser et al., 2022 |
| | | Trial times | Increased | |
| $Ca_v$1.2 KO [129SvEv] | $Ca^{2+}$ signaling | No. of lower rung touches | Decreased | Klomp et al., 2022 |
| | | Step size | No effect | |
| | | Step time | Increased | |
| $Ca_v$1.3 KO [C57BL/6NTac] | $Ca^{2+}$ signaling | No. of lower rung touches | No effect | Lauffer et al., 2022 |
| | | Step size | Decreased | |
| *Nf1*$^{+/-}$ [C57BL/6 x 129T2/SvEms] | Neuro-fibromatosis type 1 | Trials without lower rung touches | No effect | Ottenhoff et al., 2022 |
| WT [C57BL/6N] | Glial cell proliferation | | [4] | Fang et al., 2023 |
| *Pcp2-Ercc1*$^{\Delta/f}$ [C57BL/6J] | Cerebellar degeneration | No. of lower rung touches | Increased | Birkisdóttir et al., 2023 |

In this Table, we have included all peer-reviewed studies using unperturbed trials, i.e., trials in the absence of fixed or appearing obstacles, that could be retrieved via Google Scholar on August 29$^{th}$ 2023, using the query (Erasmusladder OR "Erasmus ladder" OR "Erasmus automated ladder"). The purpose of each study in ultimately understanding fundamental (white background) or clinical (yellow background) questions is indicated in the column "Model for". [1] A trial was defined as successful if the mice were able to walk on the ladder without disruption (twisting, turning, walking backwards, etc.). [2] NL3 expression specifically in Purkinje cells. [3] Decrease in steps with step size 2 during training correlated positively with the occurrence of axonal swellings of Purkinje cells in lobule III. [4] Training on the ErasmusLadder induced proliferation of oligodendrocyte precursor cells in the hippocampus. FXTAS = fragile x tremor/ataxia syndrome; HGPPS = horizontal gaze palsy with progressive scoliosis; MDCMC = congenital muscular dystrophy with megaconial myopathy.

**Table S2 – Weights (Fig. 1A)**

| Age (weeks) | Weight control (median [IQR]) | Weight PC-KO (median [IQR]) | Mann-Whitney tests | | |
|---|---|---|---|---|---|
| | | | *p* | U | Significant?[1] |
| 13 | 27.5 [2.3] g | 26.5 [1.0] g | 0.2447 | 17.5 | No |
| 14 | 30.2 [1.9] g | 29.5 [2.8] g | 0.6022 | 23.0 | No |
| 15 | 30.4 [2.4] g | 29.3 [3.1] g | 0.4871 | 21.5 | No |
| 16 | 30.7 [3.5] g | 30.5 [2.8] g | 0.6022 | 23.0 | No |
| 17 | 30.7 [4.5] g | 30.0 [2.9] g | 0.9076 | 26.5 | No |
| 18 | 30.7 [4.3] g | 30.9 [1.2] g | 0.9538 | 27.0 | No |
| 19 | 31.9 [4.6] g | 32.2 [2.1] g | 0.8665 | 26.0 | No |
| 20 | 32.2 [4.9] g | 31.8 [2.1] g | 0.9078 | 26.5 | No |
| 21 | 32.6 [4.4] g | 32.5 [2.0] g | 0.7789 | 25.0 | No |
| 22 | 32.4 [3.9] g | 32.5 [2.0] g | 0.6852 | 24.0 | No |
| 23 | 32.4 [3.0] g | 32.5 [2.0] g | 0.8166 | 25.5 | No |
| 24 | 31.9 [3.0] g | 32.1 [1.6] g | 0.9077 | 26.5 | No |
| 25 | 33.0 [3.6] g | 32.2 [1.7] g | 0.2023 | 16.5 | No |
| 26 | 34.3 [2.0] g | 32.7 [2.5] g | 0.0427 | 10.0 | No |

[1] After Benjamini-Hochberg correction for multiple comparisons.
Repeated measures ANOVA: Between subjects effect: Genotype: $F$ = 0.506, $p$ = 0.490

**Table S3 – Rotarod [2-40 rpm]: latency to fall (Fig. 1B)**

| Age (weeks) | Latency control (median [IQR]) | Latency PC-KO (median [IQR]) | Mann-Whitney tests | | |
|---|---|---|---|---|---|
| | | | *p* | U | Significant?[1] |
| 13 | [illegible] | [illegible] | [illegible] | [illegible] | No |
| 14 | [illegible] | [illegible] | [illegible] | [illegible] | No |
| 15 | [illegible] | [illegible] | [illegible] | 27.0 | No |
| 16 | [illegible] | [illegible] | [illegible] | [illegible] | No |
| 17 | [illegible] | [illegible] | [illegible] | [illegible] | No |
| 18 | [illegible] | [illegible] | [illegible] | [illegible] | No |
| 19 | [illegible] | [illegible] | [illegible] | [illegible] | No |
| 20 | [illegible] | [illegible] | [illegible] | [illegible] | No |
| 21 | [illegible] | [illegible] | [illegible] | [illegible] | No |
| 22 | [illegible] | [illegible] | [illegible] | [illegible] | No |
| 23 | [illegible] | [illegible] | [illegible] | [illegible] | No |
| 24 | [illegible] | [illegible] | [illegible] | [illegible] | [illegible] |
| 25 | [illegible] | [illegible] | [illegible] | [illegible] | [illegible] |
| 26 | [illegible] | [illegible] | [illegible] | [illegible] | No |

[1] After Benjamini-Hochberg correction for multiple comparisons.
Repeated measures ANOVA: Between subjects effect: Genotype: $F = 1.917$, $p = 0.189$

**Table S4 – Rotarod [2-80 rpm]: latency to fall (Fig. 1C)**

| Age (weeks) | Latency control (median [IQR]) | Latency PC-KO (median [IQR]) | Mann-Whitney tests | | |
|---|---|---|---|---|---|
| | | | *p* | U | Significant?[1] |
| 13 | 132 [31] s | 146 [38] s | 0.8168 | 25.5 | No |
| 14 | 162 [36] s | 158 [42] s | 0.8665 | 26.0 | No |
| 15 | 171 [17] s | 161 [47] s | 0.9538 | 27.0 | No |
| 16 | 151 [46] s | 175 [29] s | 0.1520 | 15.0 | No |
| 17 | 167 [55] s | 164 [22] s | 0.9551 | 27.0 | No |
| 18 | 153 [47] s | 157 [24] s | 0.6943 | 24.0 | No |
| 19 | 150 [68] s | 159 [13] s | 0.7789 | 25.0 | No |
| 20 | 160 [60] s | 140 [28] s | 0.1893 | 16.0 | No |
| 21 | 167 [76] s | 136 [33] s | 0.4634 | 21.0 | No |
| 22 | 150 [48] s | 120 [11] s | 0.0422 | 10.0 | No |
| 23 | 156 [79] s | 108 [23] s | 0.0727 | 12.0 | No |
| 24 | 119 [85] s | 107 [22] s | 0.7282 | 24.5 | No |
| 25 | 170 [85] s | 118 [22] s | 0.0401 | 10.0 | No |
| 26 | 161 [82] s | 98 [19] s | 0.0091 | 5.0 | No |

[1] After Benjamini-Hochberg correction for multiple comparisons.
Repeated measures ANOVA: Between subjects effect: Genotype: $F = 1.231$, $p = 0.287$

**Table S5 – Trial times PC-KO mice (Fig. 2C)**

| Age (weeks) | Trial time control (median [IQR]) | Trial time PC-KO (median [IQR]) | Mann-Whitney tests | | |
|---|---|---|---|---|---|
| | | | *p* | U | Significant?[1] |
| 13 | 1.17 [0.41] s | 1.17 [0.32] s | *0.7984* | 29.0 | No |
| 14 | 1.23 [0.34] s | 1.16 [0.33] s | *0.8785* | 30.0 | No |
| 15 | 1.17 [0.15] s | 1.19 [0.34] s | *0.5054* | 25.0 | No |
| 16 | 1.14 [0.19] s | 1.33 [0.23] s | *0.0650* | 14.0 | No |
| 17 | 1.21 [0.21] s | 1.26 [0.15] s | *0.2345* | 20.0 | No |
| 18 | 1.08 [0.19] s | 1.32 [0.38] s | ***0.0070*** | 7.0 | **Yes** |
| 19 | 1.17 [0.19] s | 1.80 [0.43] s | ***0.0003*** | 1.0 | **Yes** |
| 20 | 1.23 [0.41] s | 1.68 [0.30] s | ***0.0030*** | 5.0 | **Yes** |
| 21 | 1.06 [0.26] s | 1.87 [0.58] s | ***0.0011*** | 3.0 | **Yes** |
| 22 | 1.16 [0.16] s | 2.02 [0.44] s | ***0.0002*** | 0.0 | **Yes** |
| 23 | 1.12 [0.06] s | 2.28 [0.49] s | ***0.0002*** | 0.0 | **Yes** |
| 24 | 1.22 [0.22] s | 2.57 [0.35] s | ***0.0002*** | 0.0 | **Yes** |
| 25 | 1.21 [0.13] s | 2.40 [0.46] s | ***0.0002*** | 0.0 | **Yes** |
| 26 | 1.15 [0.19] s | 2.58 [0.35] s | ***0.0002*** | 0.0 | **Yes** |

[1] After Benjamini-Hochberg correction for multiple comparisons.

**Table S6 – Touch times PC-KO mice (Fig. 2D)**

| Age (weeks) | Touch time control (median [IQR]) | Touch time PC-KO (median [IQR]) | Mann-Whitney tests | | |
|---|---|---|---|---|---|
| | | | *p* | U | Significant?[1] |
| 13 | 84.5 [15.3] ms | 79.0 [13.0] ms | *0.5630* | *26.0* | No |
| 14 | 89.0 [12.8] ms | 81.5 [9.0] ms | *0.4933* | *25.0* | No |
| 15 | 81.5 [20.3] ms | 83.0 [12.8] ms | *0.7926* | *29.0* | No |
| 16 | 84.5 [16.0] ms | 86.5 [4.8] ms | *0.6338* | *27.0* | No |
| 17 | 85.5 [11.0] ms | 90.5 [7.8] ms | *0.3707* | *23.0* | No |
| 18 | 80.0 [12.9] ms | 100.5 [9.8] ms | ***0.0134*** | *8.0* | **Yes** |
| 19 | 93.3 [13.5] ms | 130.5 [28.3] ms | ***0.0027*** | *3.0* | **Yes** |
| 20 | 89.0 [16.5] ms | 117.0 [19.3] ms | ***0.0133*** | *8.0* | **Yes** |
| 21 | 82.5 [21.8] ms | 121.5 [29.5] ms | ***0.0039*** | *4.0* | **Yes** |
| 22 | 91.0 [19.6] ms | 130.5 [11.0] ms | ***0.0009*** | *0.0* | **Yes** |
| 23 | 86.0 [18.3] ms | 144.0 [31.1] ms | ***0.0009*** | *0.0* | **Yes** |
| 24 | 89.5 [19.1] ms | 137.5 [11.8] ms | ***0.0009*** | *0.0* | **Yes** |
| 25 | 83.0 [13.3] ms | 135.3 [28.6] ms | ***0.0009*** | *0.0* | **Yes** |
| 26 | 86.0 [10.3] ms | 137.0 [12.3] ms | ***0.0009*** | *0.0* | **Yes** |

[1] After Benjamini-Hochberg correction for multiple comparisons.

**Table S7 – Lower rung touches PC-KO mice (Fig. 2E)**

| Age (weeks) | No. of touches control (median [IQR]) | No. of touches PC-KO (median [IQR]) | Mann-Whitney tests | | |
|---|---|---|---|---|---|
| | | | *p* | U | Significant?[1] |
| 13 | 2.98 [1.78] % | 2.92 [1.70] % | *0.6363* | *27.0* | No |
| 14 | 2.60 [1.63] % | 3.33 [0.79] % | *0.4946* | *25.0* | No |
| 15 | 3.13 [1.22] % | 4.54 [1.16] % | ***0.0104*** | *8.0* | **Yes** |
| 16 | 1.71 [2.00] % | 4.16 [2.08] % | ***0.0281*** | *11.0* | **Yes** |
| 17 | 3.63 [2.68] % | 4.14 [1.76] % | *0.8785* | *30.0* | No |
| 18 | 2.28 [0.81] % | 4.46 [2.36] % | ***0.0207*** | *10.0* | **Yes** |
| 19 | 3.24 [1.21] % | 6.36 [2.93] % | ***0.0207*** | *10.0* | **Yes** |
| 20 | 2.69 [1.27] % | 6.68 [3.18] % | ***0.0003*** | *1.0* | **Yes** |
| 21 | 3.14 [1.14] % | 11.35 [3.21] % | ***0.0002*** | *0.0* | **Yes** |
| 22 | 1.78 [2.26] % | 13.66 [3.82] % | ***0.0002*** | *0.0* | **Yes** |
| 23 | 2.04 [1.91] % | 12.55 [2.51] % | ***0.0002*** | *0.0* | **Yes** |
| 24 | 2.50 [1.55] % | 15.51 [1.63] % | ***0.0002*** | *0.0* | **Yes** |
| 25 | 3.57 [2.35] % | 19.46 [4.71] % | ***0.0002*** | *0.0* | **Yes** |
| 26 | 1.91 [1.25] % | 17.17 [6.17] % | ***0.0002*** | *0.0* | **Yes** |

[1] After Benjamini-Hochberg correction for multiple comparisons.

**Table S8 – Diagonal stance PC-KO mice (Fig. 3C)**

| Age (weeks) | Time control (median [IQR]) | Time PC-KO (median [IQR]) | Mann-Whitney tests | | |
|---|---|---|---|---|---|
| | | | *p* | U | Significant?[1] |
| 13 | 31.95 [5.53] % | 28.62 [2.56] % | *0.1049* | 16.0 | No |
| 14 | 31.36 [8.39] % | 29.92 [2.60] % | *0.5737* | 26.0 | No |
| 15 | 33.17 [8.03] % | 27.81 [3.29] % | *0.1034* | 16.0 | No |
| 16 | 32.33 [6.63] % | 28.77 [2.23] % | *0.0650* | 14.0 | No |
| 17 | 31.75 [7.84] % | 29.71 [6.03] % | *0.2345* | 20.0 | No |
| 18 | 30.91 [5.28] % | 27.99 [7.02] % | *0.2786* | 21.0 | No |
| 19 | 28.89 [5.53] % | 28.73 [1.79] % | *0.7984* | 29.0 | No |
| 20 | 29.83 [6.44] % | 30.54 [1.81] % | *0.4418* | 24.0 | No |
| 21 | 26.64 [9.54] % | 29.17 [6.00] % | *0.9591* | 31.0 | No |
| 22 | 30.54 [9.31] % | 28.62 [7.02] % | *0.5737* | 26.0 | No |
| 23 | 29.05 [4.80] % | 29.57 [2.76] % | *0.9591* | 31.0 | No |
| 24 | 28.08 [6.32] % | 24.36 [4.05] % | *0.0148* | 9.0 | No |
| 25 | 26.41 [5.67] % | 23.23 [4.85] % | *0.0379* | 12.0 | No |
| 26 | 27.83 [5.21] % | 22.55 [5.23] % | *0.0207* | 10.0 | No |

[1] After Benjamini-Hochberg correction for multiple comparisons.

**Table S9 – Trial times PC-Sca1 mice (Fig. 4D)**

| Age (weeks) | Trial time control (median [IQR]) | Trial time PC-Sca1 (median [IQR]) | Mann-Whitney tests | | |
|---|---|---|---|---|---|
| | | | *p* | U | Significant?[1] |
| 7 | 1.74 [0.83] s | 1.88 [0.59] s | *0.5619* | 51.0 | No |
| 8 | 1.67 [0.53] s | 1.76 [0.31] s | *0.5619* | 51.0 | No |
| 9 | 1.49 [0.44] s | 1.80 [0.63] s | *0.0557* | 31.0 | No |
| 10 | 1.41 [0.37] s | 1.73 [0.53] s | *0.0336* | 28.0 | No |
| 11 | 1.67 [0.51] s | 1.72 [1.73] s | *0.3000* | 44.0 | No |
| 12 | 1.83 [0.76] s | 1.90 [1.61] s | *0.6994* | 54.0 | No |
| 13 | 1.73 [0.59] s | 1.62 [1.01] s | *0.6522* | 53.0 | No |
| 14 | 1.85 [1.08] s | 1.74 [1.25] s | *0.7477* | 55.0 | No |
| 15 | 1.37 [0.37] s | 2.22 [1.25] s | *0.1014* | 35.0 | No |
| 16 | 1.70 [0.61] s | 2.06 [1.45] s | *0.4779* | 49.0 | No |
| 17 | 1.62 [0.55] s | 2.63 [2.04] s | *0.0165* | 23.5 | No |
| 18 | 1.44 [0.51] s | 2.45 [1.46] s | *0.0233* | 26.0 | No |
| 19 | 1.53 [0.79] s | 3.70 [2.41] s | *0.0158* | 24.0 | No |
| 20 | 1.43 [0.54] s | 2.80 [2.06] s | ***0.0032*** | 17.0 | **Yes** |

[1] After Benjamini-Hochberg correction for multiple comparisons.

**Table S10 – Touch times PC-Sca1 mice (Fig. 4E)**

| Age (weeks) | Touch time control (median [IQR]) | Touch time PC-Sca1 (median [IQR]) | Mann-Whitney tests | | |
|---|---|---|---|---|---|
| | | | *p* | U | Significant?[1] |
| 7 | 94.0 [25.5] ms | 106.0 [17.3] ms | *0.1228* | 36.5 | No |
| 8 | 94.0 [14.5] ms | 106.0 [20.3] ms | *0.1149* | 36.0 | No |
| 9 | 80.5 [23.5] ms | 117.0 [22.0] ms | *0.0138* | 22.5 | No |
| 10 | 95.0 [19.3] ms | 108.0 [13.3] ms | *0.0281* | 27.0 | No |
| 11 | 94.5 [19.5] ms | 100.0 [26.0] ms | *0.3574* | 46.0 | No |
| 12 | 94.0 [14.8] ms | 103.0 [22.0] ms | *0.2238* | 41.5 | No |
| 13 | 86.0 [15.0] ms | 96.0 [18.3] ms | *0.1886* | 40.0 | No |
| 14 | 90.0 [9.5] ms | 97.5 [24.3] ms | *0.1308* | 37.0 | No |
| 15 | 90.0 [13.8] ms | 90.0 [21.5] ms | *0.4302* | 48.0 | No |
| 16 | 91.0 [15.3] ms | 104.0 [15.3] ms | *0.0613* | 31.5 | No |
| 17 | 93.0 [22.5] ms | 112.5 [39.5] ms | *0.0417* | 29.0 | No |
| 18 | 95.0 [15.0] ms | 112.0 [15.3] ms | *0.0302* | 27.0 | No |
| 19 | 91.0 [16.0] ms | 129.0 [26.8] ms | *0.0095* | 20.5 | No |
| 20 | 89.0 [12.8] ms | 118.0 [41.0] ms | ***0.0012*** | 10.5 | **Yes** |

[1] After Benjamini-Hochberg correction for multiple comparisons.

**Table S11 – Lower rung touches PC-Sca1 mice (Fig. 4G)**

| Age (weeks) | No. of touches control (median [IQR]) | No. of touches PC-Sca1 (median [IQR]) | Mann-Whitney tests | | |
|---|---|---|---|---|---|
| | | | *p* | U | Significant?[1] |
| 7 | 4.46 [0.66] % | 7.55 [6.37] % | ***0.0128*** | 23.0 | **Yes** |
| 8 | 4.79 [3.37] % | 7.47 [5.93] % | *0.4779* | 49.0 | No |
| 9 | 4.03 [0.26] % | 7.63 [3.44] % | ***0.0008*** | 12.0 | **Yes** |
| 10 | 6.32 [1.02] % | 10.13 [2.50] % | ***0.0052*** | 19.0 | **Yes** |
| 11 | 4.49 [0.87] % | 10.92 [3.61] % | ***0.0001*** | 5.0 | **Yes** |
| 12 | 6.16 [0.93] % | 8.11 [2.62] % | *0.2703* | 43.0 | No |
| 13 | 3.93 [2.83] % | 9.47 [5.27] % | ***0.0019*** | 15.0 | **Yes** |
| 14 | 4.69 [1.16] % | 9.52 [5.30] % | ***0.0003*** | 9.0 | **Yes** |
| 15 | 4.35 [1.10] % | 11.43 [4.90] % | ***0.0001*** | 2.0 | **Yes** |
| 16 | 4.38 [0.48] % | 9.88 [3.86] % | ***0.0001*** | 5.0 | **Yes** |
| 17 | 4.69 [1.90] % | 13.08 [4.97] % | ***0.0001*** | 0.0 | **Yes** |
| 18 | 4.55 [0.61] % | 11.38 [3.93] % | ***0.0001*** | 0.0 | **Yes** |
| 19 | 3.26 [0.98] % | 11.38 [1.93] % | ***0.0001*** | 1.0 | **Yes** |
| 20 | 3.31 [2.41] % | 11.38 [2.47] % | ***0.0004*** | 10.0 | **Yes** |

[1] After Benjamini-Hochberg correction for multiple comparisons.

**Table S12 – Large steps PC-Sca1 mice (Fig. 4H)**

| Age (weeks) | No. of steps control (median [IQR]) | No. of steps PC-Sca1 (median [IQR]) | Mann-Whitney tests | | |
|---|---|---|---|---|---|
| | | | *p* | U | Significant?[1] |
| 7 | 73.89 [7.32] % | 51.21 [27.55] % | *0.0652* | 32.0 | No |
| 8 | 70.59 [6.95] % | 63.24 [18.28] % | *0.3000* | 44.0 | No |
| 9 | 77.19 [4.33] % | 61.18 [16.89] % | ***0.0066*** | 20.0 | **Yes** |
| 10 | 70.86 [3.28] % | 63.92 [13.60] % | ***0.0010*** | 13.0 | **Yes** |
| 11 | 76.92 [4.62] % | 59.91 [20.05] % | ***0.0024*** | 16.0 | **Yes** |
| 12 | 69.66 [6.34] % | 63.64 [30.40] % | *0.1014* | 35.0 | No |
| 13 | 77.74 [3.39] % | 59.46 [24.81] % | *0.0879* | 34.0 | No |
| 14 | 74.62 [11.92] % | 59.78 [26.67] % | *0.0473* | 30.0 | No |
| 15 | 75.96 [3.05] % | 55.23 [22.78] % | ***0.0128*** | 23.0 | **Yes** |
| 16 | 73.91 [7.52] % | 57.63 [40.16] % | ***0.0010*** | 13.0 | **Yes** |
| 17 | 72.46 [6.27] % | 50.00 [32.35] % | ***0.0024*** | 16.0 | **Yes** |
| 18 | 77.92 [5.69] % | 51.69 [43.53] % | ***0.0066*** | 20.0 | **Yes** |
| 19 | 77.95 [8.80] % | 51.41 [28.61] % | ***0.0010*** | 13.0 | **Yes** |
| 20 | 76.97 [7.74] % | 53.53 [31.33] % | ***0.0003*** | 9.0 | **Yes** |

[1] After Benjamini-Hochberg correction for multiple comparisons.
